# State Dependent Coupling of Hippocampal Oscillations

**DOI:** 10.1101/2022.05.03.490415

**Authors:** Brijesh Modi, Matteo Guardamagna, Federico Stella, Marilena Griguoli, Enrico Cherubini, Francesco P. Battaglia

## Abstract

Oscillations occurring simultaneously in a given area represent a physiological unit of brain states. They allow for temporal segmentation of spikes and support distinct behaviors. To establish how multiple oscillatory components co-varies simultaneously and influence neuronal firing during sleep and wakefulness, we describe a multi-variate analytical framework for constructing the state space of hippocampal oscillations. Examining the co-occurrence patterns of oscillations on the state space, across species, uncovered the presence of network constraints and distinct set of cross-frequency interactions during wakefulness as compared to sleep. We demonstrated how the state space can be used as a canvas to map the neural firing and found that distinct neurons during navigation were tuned to different set of simultaneously occurring oscillations during sleep. This multivariate analytical framework provides a window to move beyond classical bivariate pipelines, for investigating oscillations and neuronal firing, thereby allowing to factor-in the complexity of oscillation-population interactions.

## INTRODUCTION

Oscillations, generated by large neuronal ensembles, are a hallmark of the mammalian brain. They are well preserved during evolution (Buzsáki and Draguhn, 2004) and have been suggested to play a key role in high cognitive functions (Uhlhaas and Singer, 2010). They enable the synchronization of neural activity within and across brain regions, thus promoting the precise temporal coordination of neural processes underlying memory, perception, and behavior (Colgin and Moser, 2010). Their disruption leads to cognitive and sensorimotor deficits associated with several neuropsychiatric diseases (Uhlhaas et al., 2011). Oscillations occurring at different frequency bands (from 0.5 to 200 Hz), interact with each other through hierarchical cross-frequency coupling (González et al., 2020; Jensen and Colgin, 2007; Tort et al., 2008). Such interactions are linked to different computational operations, or different phases in a computational operation and are thought to characterize distinct brain states supporting various aspects of behavior (McCormick et al., 2020).

In the hippocampus, theta cycles are often nested with bouts of faster oscillations in the gamma frequency and are thought to be related to different stages of memory processing (Lopes-Dos-Santos et al., 2018). Specifically, during wakefulness, the CA1 region is characterized by cross-frequency coupling between theta and distinct gamma oscillations which exhibit their peak power at distinct phases of ongoing slow oscillations (Belluscio et al., 2012; Buzsáki and Wang, 2012; Colgin et al., 2009; Csicsvari et al., 2003; Scheffer-Teixeira et al., 2012; Yamamoto et al., 2014). The slow, medium and fast gamma oscillations, emerging from *stratum radiatum (SR), stratum lacunosum moleculare (SLM)* and *stratum pyramidale (SP)* respectively, are linked to network activity in entorhinal cortex, CA3 and CA1 region respectively (Colgin et al., 2009; Fernández-Ruiz et al., 2017; Guardamagna et al., 2021; Lasztóczi and Klausberger, 2016; Schomburg et al., 2014). During sleep, theta and medium gamma oscillations dominate the REM sleep, whereas delta, spindle frequency band (11-20 Hz, hereafter spindle) and ripples or fast gamma (100-200 Hz), linked to memory consolidation processes, dominate the non-REM sleep (Battaglia et al., 2011; Buzsáki, 1989; Genzel et al., 2014; Roumis and Frank, 2015). The composition of hippocampal states by distinct set of slow and fast oscillations reflects synchrony among different brain areas and corresponds to distinct functional states of the hippocampal network (Colgin, 2015). However, despite extensive research on simultaneously occurring oscillations, very little is known about the dynamic variations in the composition of hippocampal network state with time and behavior.

Oscillations are generated by neuronal population, and they in turn, are known to modulate the firing activity of individual cells (Benchenane et al., 2010; Hulse et al., 2017; O’Keefe and Recce, 1993). Classically, neuronal firing has been characterized in the context of a single frequency band during distinct brain states (Buzsáki and Draguhn, 2004; Fox et al., 1986; Grosmark et al., 2012; Hafting et al., 2008; Henze et al., 2000; Siapas et al., 2005; Wingerden et al., 2010). However, the classical analytical methods, due to their bivariate nature, are limited to examining the firing activity of neurons in relation to power or phase or frequency of a specific oscillation only, thus neglecting how multiple oscillations, occurring simultaneously in a given region, modulate the neuronal firing in a combinatorial manner.

Here, a novel analytical approach has been described to investigate how simultaneously occurring network oscillations dynamically contributes to the composition of hippocampal states and how they influence the neuronal firing. To this aim, multisite local field potentials (LFPs), recorded from the CA1 region of the dorsal hippocampus of freely moving mice, were used to construct the network state space during wakefulness and sleep. This method has allowed to create a compact representation of the state of multiple oscillatory processes, distinct in frequency and anatomical localization. We used this compact representation for studying various oscillations, their intrinsic organization, their temporal progression and their simultaneous influence on distinct neuronal ensembles and behavior. In addition, the state space provided a window for observing the hippocampal population in the context of the network in which they are embedded. This allowed us to examine how cell fires as a function of the network state and how multiple oscillations simultaneously modulate the cell firing. As a proof of concept, this approach has been applied to datasets recorded, in the similar experimental conditions, from another animal species (rats), in Dr. Gyorgy Buzsaki’s lab at NYU (*hc-11 dataset, crncs.org* (Grosmark and Buzsáki, 2016; Grosmark et al., 2016)). Lastly, the network state space framework has been applied to study alterations in the organization of hippocampal oscillations in mice lacking neuroligin 3 (NLG3 KO; neuroligin-3 knock out), an animal model of autism (Baudouin et al., 2012).

## RESULTS

### Experimental Paradigm and Construction of the Network State Space

On a single day, we recorded brain activity in freely moving mice (n=4) during 4 consecutive trials: (1) Sleep1, (2) Novel Arena-1 Exploration, (3) Novel Arena-2 Exploration, (4) Sleep2, (Fig 1A). Layer-resolved local field potential were extracted from the CA1 region of the dorsal hippocampus using multisite silicon probes (Fig 1B; Guardamagna et al., 2022). LFPs from SP, SR and SLM layers were used to extract signals in the following frequency bands: delta (1-5 Hz), theta (6-10 Hz), spindle (10-20 Hz), slow gamma (20-45 Hz), medium gamma (60-90 Hz) and fast gamma (100-200 Hz). Each of these 18 frequency bands (6 from each layer, Fig. 1C), were used to compute median power in non-overlapping bins of 200 milliseconds and then smoothed using Gaussian kernel. The resulting 18 power time series across all 4 trials combined (divided into say, N bins of 200 milliseconds each) forms a cloud of N points in 18-dimensional space, which we refer to as the network state space (Fig 2A). Each point in the cloud represents the state of the network (i.e., the power configuration of the 18 frequency bands from 3 layers) at a given time. Uniform Manifold Approximation and Projection (UMAP, McInnes et al., 2020), was employed to reduce the dimensionality of the network state space from 18D to 2D. (Fig 1C, Fig 2A). A similar state space was constructed using the power in current source density (CSD) signals, instead of LFPs (Fig S3). We next characterized some fundamental properties of the network state space during sleep and wakefulness.

**Fig 1.**
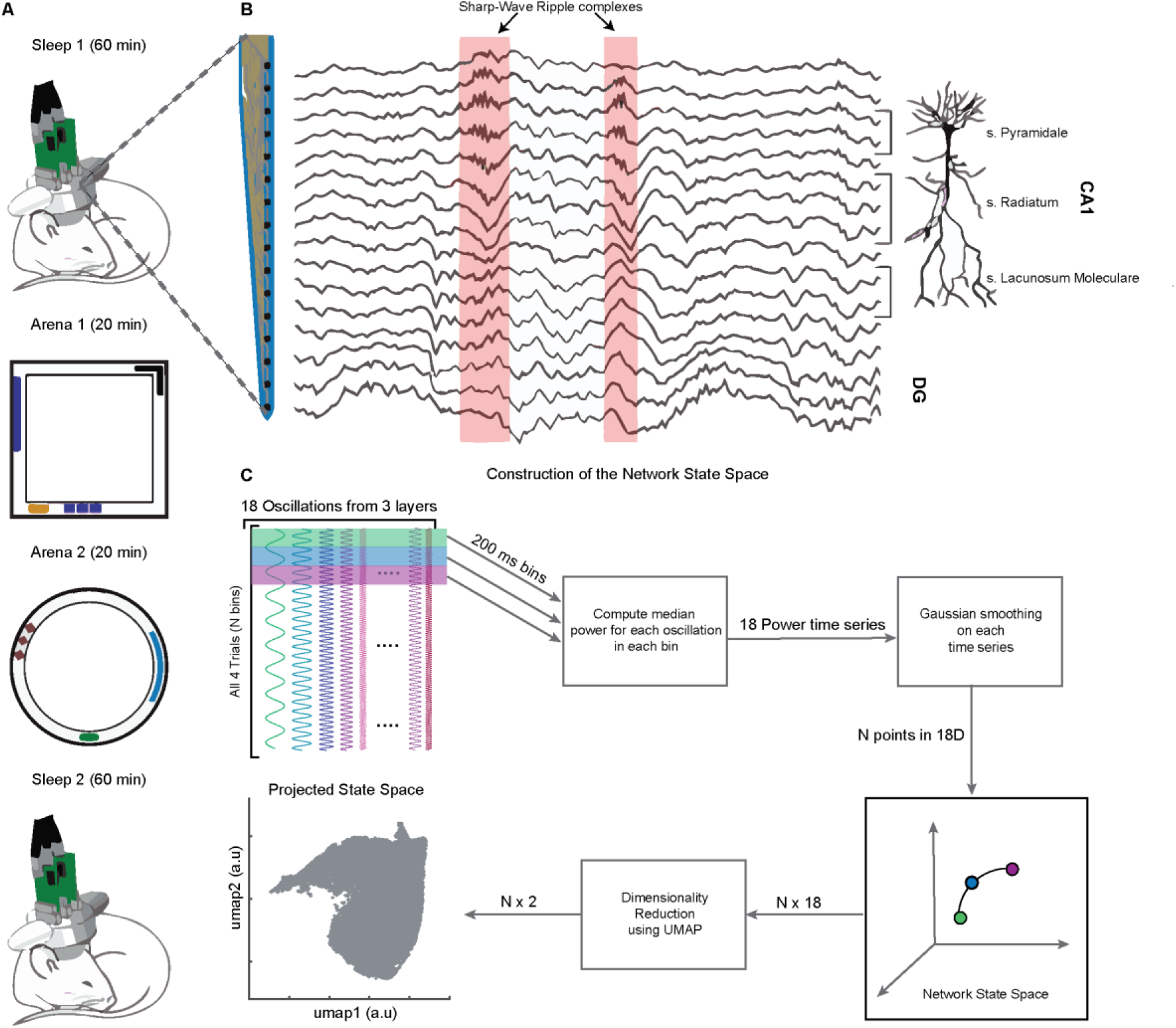
Experimental Paradigm and Pipeline for Construction of the Network State Space. A) Representative 4-trial sequence for recording during sleep and awake exploration (top to bottom). B) Representative traces of LFP recorded from various layers of dorsal CA1 using silicon probe. C) Pipeline for the construction of network state space. See also Figure S1 and S2.

**Fig 2.**
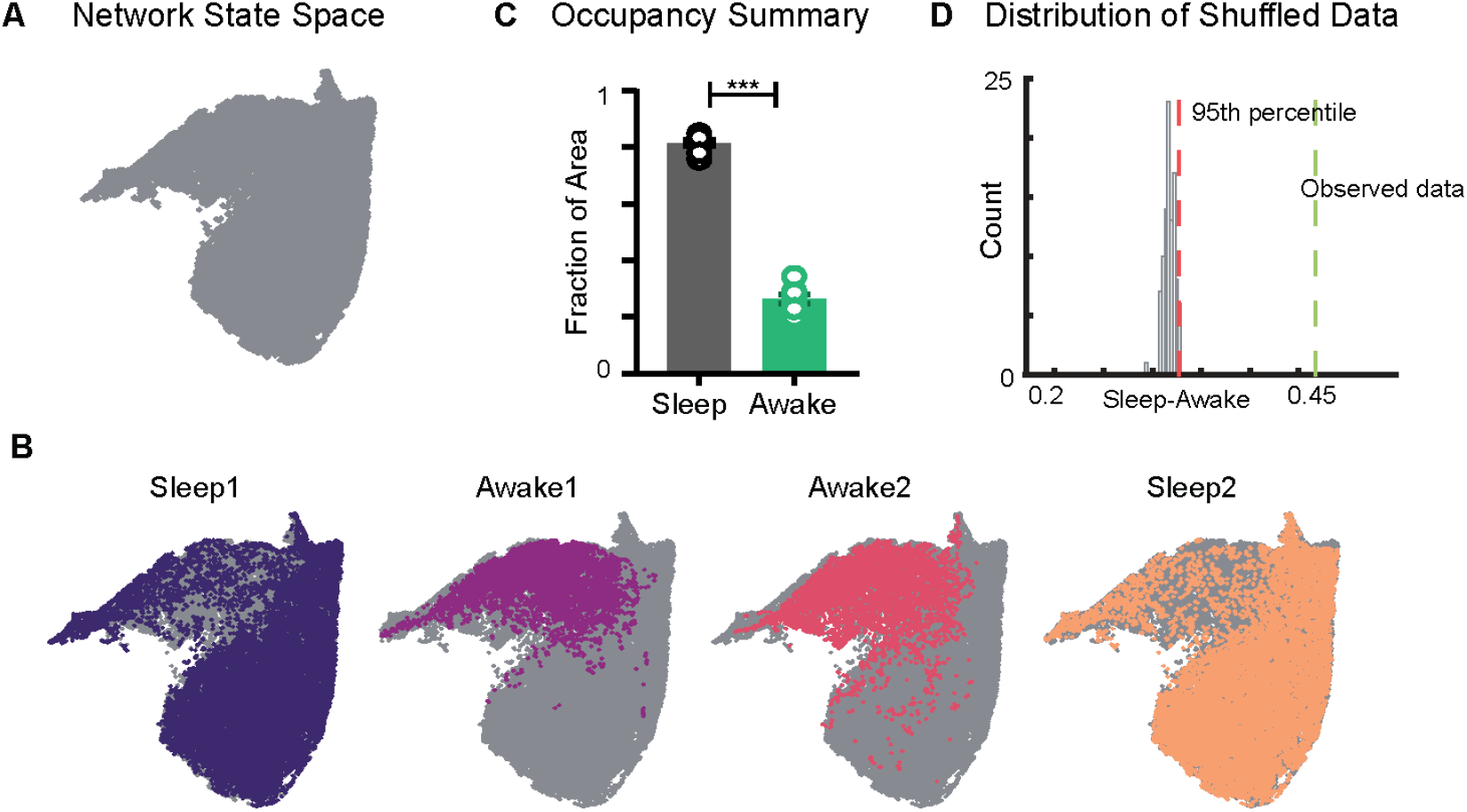
Restricted Occupancy of the Network State Space during Wakefulness. A) Network state space for all 4 trials combined. B) Trial-specific states (colored) and unvisited states (gray) on the network state space. C) Fraction of state space occupied during sleep (0.81 ± 0.01) and awake (0.26 ± 0.01) trials, suggesting significant restrictions during awake trials, (n = 8 sleep and awake trials from 4 mice. p = 0.0002, Mann-Whitney test). D) Distribution of difference in fraction of state space occupied during sleep and awake trials (*Sleep-Awake*) of surrogate data and observed data. See also Figure S3-S6.

### Restricted Occupancy of the Network State Space during Wakefulness

We first examined the occupancy of the network state space during sleep and awake exploration by overlaying trial specific states (states that network visits in a given trial) on the network state space (Fig 2A, B). We observed that during awake trials, the activity of the network was restricted to a subset of the states whereas during sleep periods, it occupied a significantly larger area of the state space. This was quantified by computing the fraction of the state space occupied during each trial. In all four mice (8 sleep and 8 awake trials), we observed a significant restriction (Fig 2C, S3) of the state space’s occupancy during wakefulness as compared to sleep (0.26 ± 0.01 *versus* 0.81 ± 0.01). The observed restriction on the state space was further compared against a surrogate data generated by randomly shuffling the binned power in all frequency bands (see Methods). We found that the observed difference in occupancy between sleep and awake trials was significantly different from those computed in surrogate data (Fig 2D). To assess whether the observed restricted occupancy was due to smaller trial duration of awake trails (20 min) as compared to sleep (60 min), we performed control analysis by computing occupancy on the state space obtained by randomly selecting equal number of network states from awake and sleep trials. We observed similar restrictions (Fig S3C) suggesting that the restricted occupancy during wakefulness is independent of trial duration. Next, we assessed whether the observed restrictions on the state space are driven by large fluctuations in low frequency bands, which may weigh preponderantly in the UMAP. We computed the state space with normalized features (Fig S4) as well as obtained the state space projection using an alternative dimensionality reduction approach (Principal Component Analysis, Fig S6). Similar patterns of restrictions on the network state space, during wakefulness, across all different projections were observed. To further validate our findings, we computed the network state space and occupancy for the datasets recorded from freely moving rats in similar experimental conditions (Fig S5) and observed remarkable similarities in restrictions on the state space, suggesting that functional organization of network oscillations during sleep and wakefulness are conserved across species. In addition, we observed that the network occupies the same set of states when animals explore distinct arenas (Fig 2B, S3). Hence, the restrictions are primarily dependent on the behavioral state of the animal rather than environmental factors.

### Characterization of Restrictions on the Network State Space during Wakefulness

To characterize the network restrictions and to visualize how different hippocampal rhythms vary simultaneously during sleep and awake trials, we overlaid the power of each oscillation on the network state space (Fig 3A). While classical analytical approaches employ bivariate methods by evaluating each oscillation individually across sleep and awake trials (Fig 3B), the network state space employ multi-variate approach and highlights how fluctuations in power for one oscillation link to the general oscillatory state of the network. This allows visualizing how power of one oscillation varies in the context of all other oscillations. In addition, it underscores the regime of operation for different oscillations during sleep and wakefulness. For instance, the restricted subspace of the state space occupied during wakefulness corresponds to states with low delta power, moderate spindle power and moderate to high power in all gamma bands. However, during sleep, the network oscillations exhibit broader range of operations since power of each oscillation varies from its minima to maxima as evident in the overlay maps (Fig 3A). This characteristic distribution during sleep was further used to determine the localization of rapid eye movement (REM), non-REM, and intermediate sleep states on the network state space.

**Fig 3.**
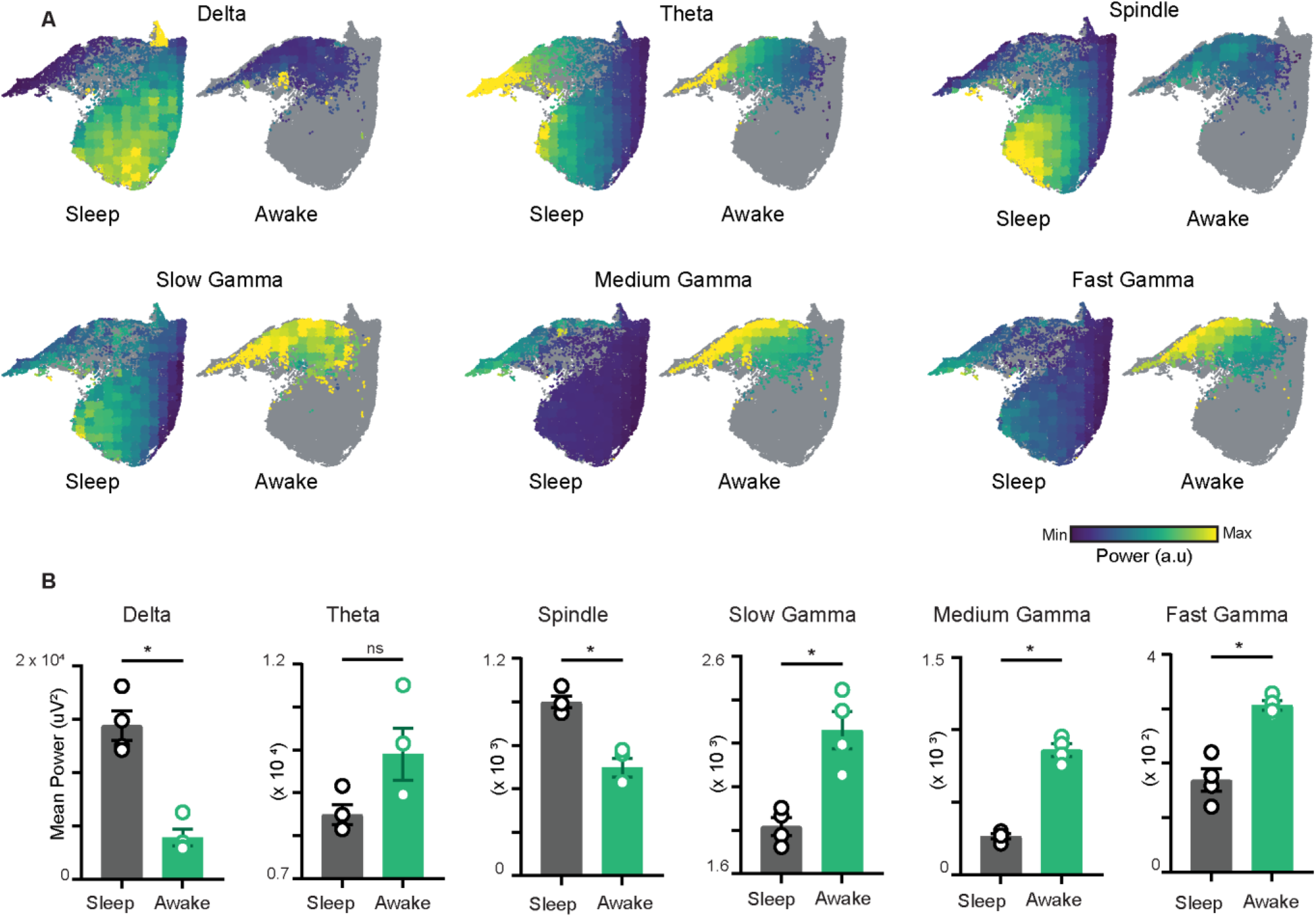
Characterization of Restrictions on the Network State Space during Wakefulness. A) Distribution of power on state space for 6 oscillations of layer stratum pyramidale: delta (1-5 Hz), theta (6-10 Hz), spindle (10-20 Hz), slow gamma (20-45 Hz), medium gamma (60-90 Hz) and fast gamma (100-200 Hz). Each oscillation is individually color-scaled to its respective minimum and maximum power. Unvisited states are colored in gray. The overlay maps demonstrate how the power of each oscillation varies on the state space and across sleep and awake trial. B) Mean power comparison between awake and sleep trials using classical approach (n = 4 mice). All p < 0.05, except for theta (p = 0.11), Mann-Whitney test.

### Distinct Localization of REM and Non-REM Sleep States on State Space

Sleep trials were further classified into REM, non-REM and Intermediate states using classical definitions (see Methods). We overlaid the *theta/delta* power ratio for each state on the network state space (Fig 4A). States with high theta/delta ratio were identified as REM (Fig 4A, C, D). Similarly, non-REM sleep states were detected by overlaying *delta* x *spindle* power product for each state on the state space (Fig 4B, C, D). The remaining states were classified as intermediate sleep states. These overlay maps allowed to visualize how functionally distinct sleep states are localized on the state space and they were further used to study how network properties varies in distinct regions of the state space. Visual inspection of power overlay maps (Fig 3A and REM and non-REM states localization maps Fig 4A, B) revealed that REM states are characterized by moderate to high power in spindle, slow, medium and fast gamma oscillatory bands simultaneously with theta oscillations whereas non-REM states are characterized by the presence of moderate to high power in theta, slow, and fast gamma oscillations along with delta and spindle. Notably, the medium gamma oscillations operate in low power mode during non-REM sleep. This co-occurrence of network oscillations is formally quantified in later sections.

**Fig 4.**
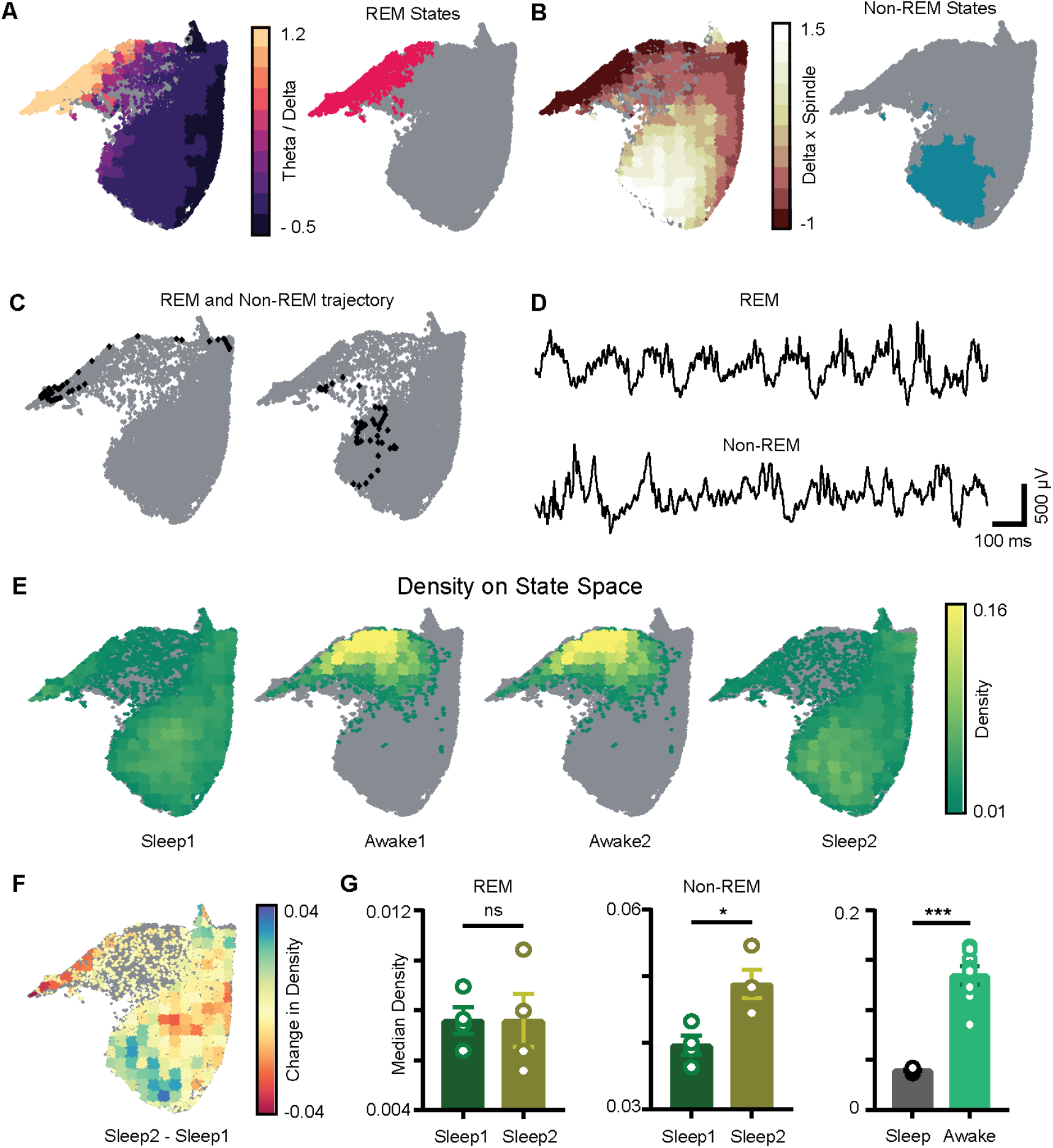
Localization of REM and Non-REM Sleep States and Density on the Network State Space. A) Left: Standardized theta/delta ratio from pyramidal layer overlaid on state space. Right: Identified REM states (magenta colored). B) Left: Standardized Delta x Spindle product from pyramidal layer overlaid on state space. Right: Identified Non-REM states (cyan colored). C) Representative REM and Non-REM states on the network state space D) Corresponding REM and Non-REM LFP from pyramidal layer. E) Density overlaid on state space across all 4 trials (unvisited states are colored gray). F) Representative change in density map across two sleep trials (sleep2 - sleep1). G) Left: Comparison of median REM density between two sleep trials (n = 4 mice); (Sleep1: 0.0076 ± 5*e* − 4 v/s Sleep2: 0.0075 ± 0.001, p = 0.88 Mann-Whitney test); Center: Comparison of median Non-REM density between two sleep trials (n = 4 mice); (Sleep1: 0.039 ± 0.001 v/s Sleep2: 0.48 ± 0.002, p < 0.05Mann-Whitney test); Right: Median density comparison between and sleep and awake trials (n = 8 sleep and awake trials, 4 mice) (0.03 ± 0.0005 v/s 0.13 ± 0.009, p = 0.0002 Mann-Whitney test)

### Increased Density of Non-REM States Associated with Theta and Gamma Oscillations after Exploration

We next computed network density which represents the fraction of time the network spends in each bin on the network state space (Fig. 4E). It is measured in the units: *number of state visits/bin/second*. Density on the state space computed during awake trials measures the normalized occurrence of corresponding behaviors during exploration whereas density computed during sleep trials measures the time spent by the network in various sleep states (REM, non-REM). Mirroring the smaller occupied state space region, the median density of awake trials was significantly higher than that detected during sleep (0.13 ± 0.009 *versus* 0.03 ± 0.0005). We next investigated the density across sleep trials (sleep1 and sleep2) by comparing the median density of REM and non-REM states between the two sleep trials (Fig 4F). We observed an increase in the median density of non-REM states during post exploration sleep (Fig 4G: center; Sleep1: 0.039 ± 0.001 *versus* Sleep2: 0.048 ± 0.002) while the density of REM sleep remained identical (Fig 4C: left; Sleep1: 0.0076 ± 5*e* − 4 *versus* Sleep2: 0.0075 ± 0.001).

Although REM and non-REM sleep states were identified using classical methods, the state space allowed quantifying changes in density of all other oscillations such as theta, slow, medium and fast gamma that occur simultaneously with delta and spindle band (from power distribution maps Fig 3A and REM and non-REM maps Fig 4A). Thus, identification of sleep states on state space provides additional information about oscillation specific changes across sleep trials. However, whether other oscillations may be statistically related to delta and spindle during non-REM states of sleep and whether such correlation depends on the state of the network remains to be elucidated.

### State Dependent Coupling of Hippocampal Oscillations

To investigate state dependent correlation among various oscillatory processes, we computed pairwise correlation matrix using binned power in all 18 frequency bands (see Methods). Each correlation matrix represents how various bands are coupled during sleep and awake trials (Fig 5A). We observed distinct configuration of frequency band’s correlations during sleep and awake trails (Fig 5B). We further investigated how coupling of bands varies on the state space by binning the state space (Fig 5C) and computed correlation matrix in each bin. We classified the bins into one of the following states: Awake, REM, non-REM, and Intermediate Sleep (see Methods). We employed UMAP to visualize the variability across correlation matrices. (Fig 5D) and observed distinct coupling configurations for awake, REM and non-REM states (Fig 5E). During REM, we found increased coupling (positive correlation) among spindle and slow gamma oscillations as well as among medium and fast gamma bands accompanied by decoupling (negative correlation) with delta. This organization among oscillations was altered during non-REM sleep as all oscillations except delta were coupled. Together these findings suggest state dependent coupling of hippocampal oscillations.

**Fig 5.**
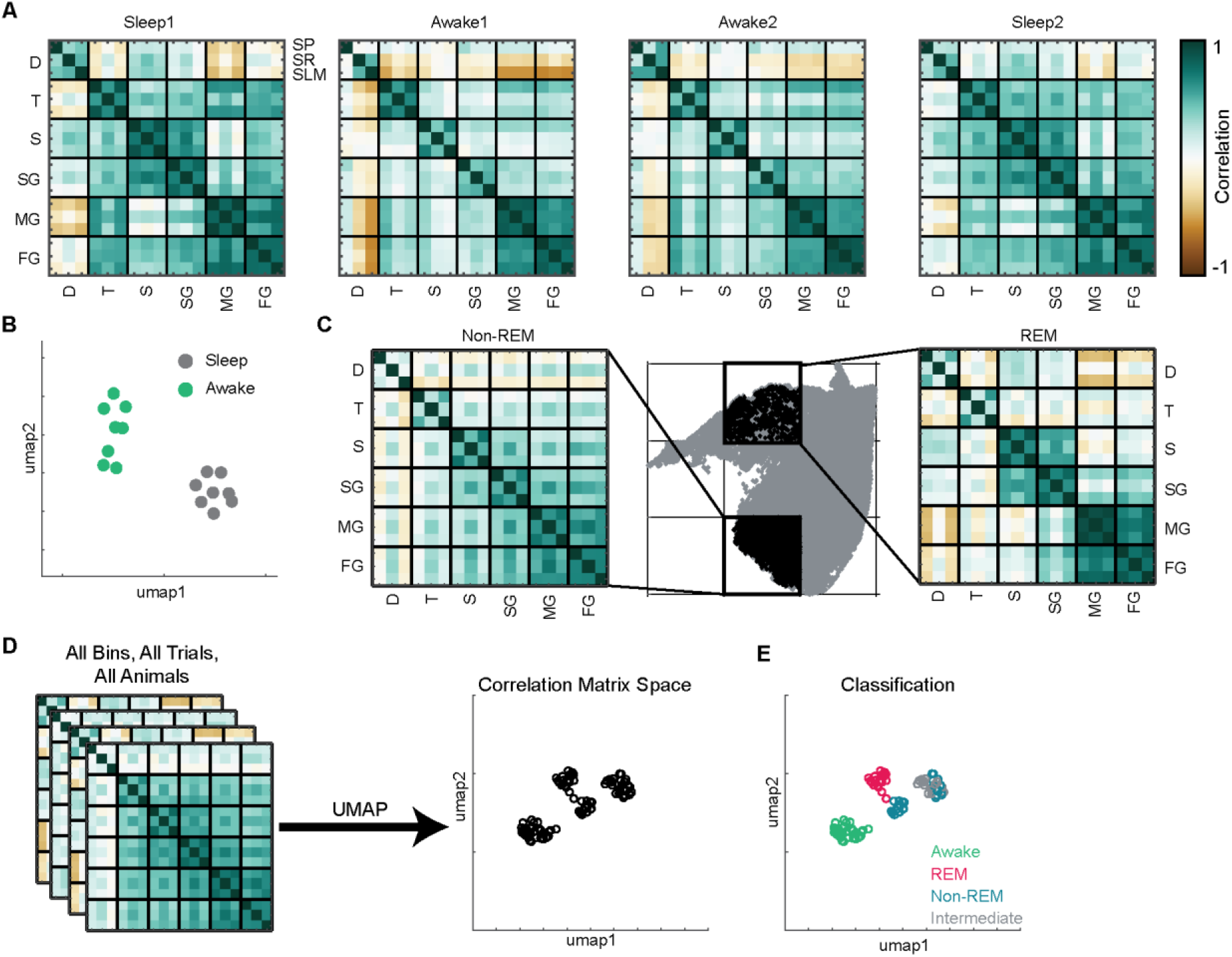
State Dependent Coupling of Hippocampal Oscillations. A) Representative pairwise correlation among 18 oscillations (D - Delta, T - Theta, S - Spindle, SG - Slow Gamma, MG - Medium Gamma, FG - Fast Gamma) from 3 layers (SP, SR, SLM). Rows are arranged by combining all 3 layers for each oscillation. B) Correlation matrix space generated by using matrices in A from all the mice (total 16 points, 8 sleep and 8 awake trials from 4 mice). Each point represents the correlation matrix of oscillations. The separation of sleep and awake points in the correlation matrix space suggests distinct nature of coupling among oscillations during sleep and wakefulness. C) Binned state space (gray) for a sleep trial. Representative REM and non-REM bins highlighted (black) and their corresponding correlation matrix of oscillations. D) Correlation matrix space constructed using matrices from all bins collected from all trials and all animals. Each point on correlation matrix space projection represents a correlation matrix. E) Bin status (Awake, REM, Non-REM, Intermediate sleep) overlaid on correlation matrix space, exhibiting state dependent coupling of hippocampal oscillations.

After characterizing the static properties of state space such as occupancy, power distribution, localization of sleep states, network state density and coupling among oscillations, in the following analyses we addressed the dynamic properties of state space such as network flow, state transitions and speed on state space.

### Plastic Nature of State Transitions on the Network State Space

To investigate how the network flows from one region of the state space to another, we plotted incoming and outgoing trajectories for all states in each bin (Fig 6A, S7, S8A). The incoming and outgoing trajectories represent the state visited by the network one second preceding or following the occurrence of a given state, respectively. They allowed visualizing the general oscillatory state of the hippocampal network varies dynamically with time. We quantified these dynamic variations on the state space by computing the transition matrix (Fig 6B, S8B, see Methods) which represents the probability by which the network switches from one state to another.

**Fig 6.**
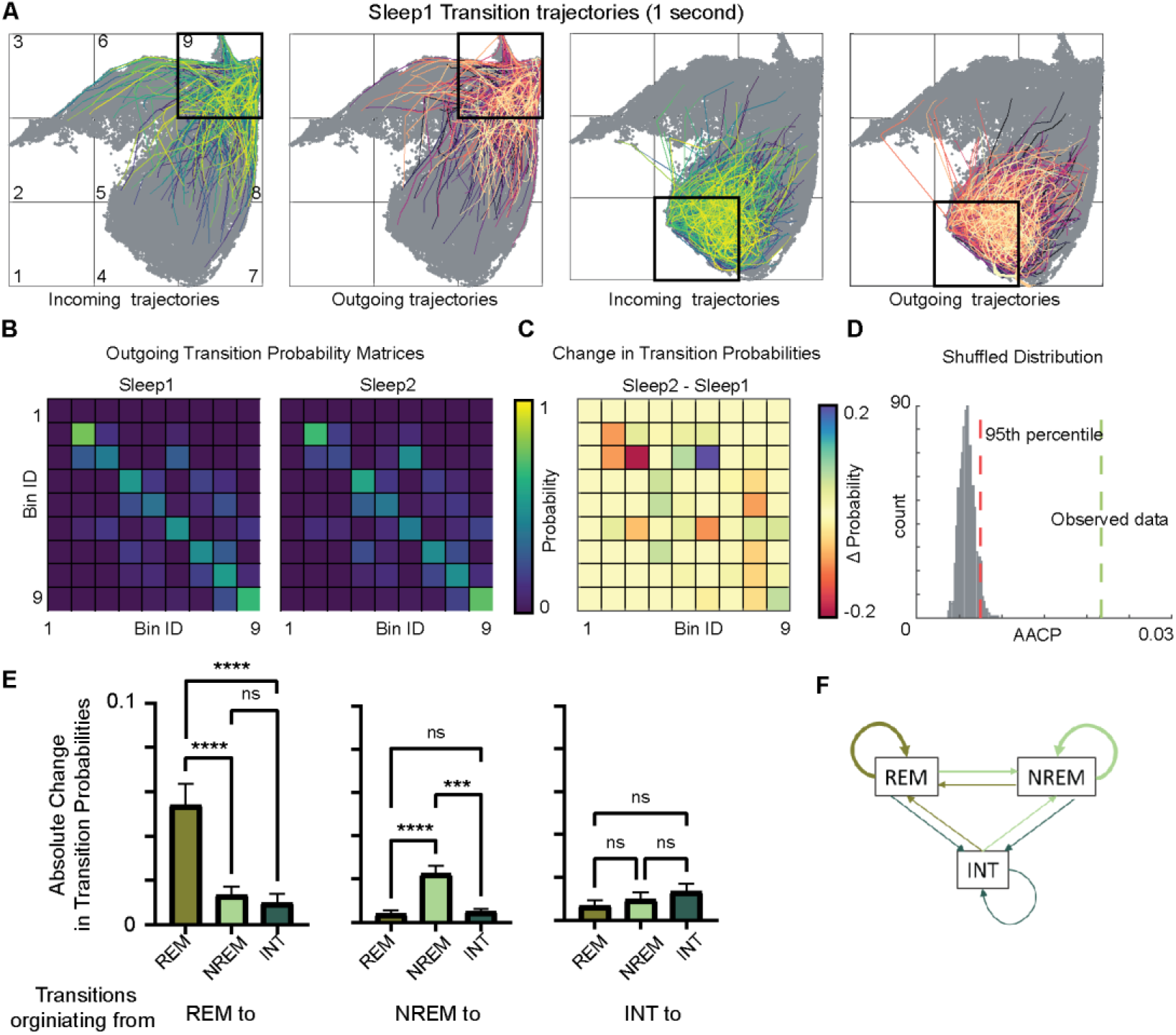
Plastic Nature of State Transitions on the Network State Space. A) Incoming and outgoing trajectories for two representative bins on state space during a sleep trial. Each trajectory spans 1 second in time before (incoming) and after (outgoing) the occurrence of given state in each bin. B) State transition matrices for two sleep trials: pre-exploration sleep (sleep1) and post-exploration sleep (sleep2). C) Change in transition probabilities across two sleep trials obtained by subtracting two transition matrices (sleep2 - sleep1). D) Comparison of observed sleep data’s average absolute change in probability with shuffled sleep data. AACP across sleep trials for observed data = 0.021, 95th percentile of shuffled data = 0.007. E) AACP across sleep trials for transitions originating from REM, non-REM and intermediate state to REM, non-REM, and intermediate sleep state. (REM: p < 0.0001, non-REM: p < 0.0001, INT: p = 0.38, one-way ANOVA) F) Schematic diagram of absolute change in transition probabilities across sleep trials. Arrow’s thickness corresponds to absolute change in transition probabilities. See also Figure S7 and S8.

We next investigated whether the paths described by the network on the state space are rigid or plastic. We computed the difference in transition matrices between two sleep or two awake trials. The resultant difference matrix (Fig 6C, S8C) highlights the bins that undergoes changes in transition probabilities. We quantified these changes in transition probabilities by computing average absolute change in probability (AACP, see Methods). This allowed to estimate how probable paths change across two sleep or two awake trials. We then assessed whether changes in transition probabilities between two sleep or two awake trails are subject to random variability or not. We generated 1000 pairs of sleep and awake trials by randomly shuffling state transitions from sleep or awake trial data and computed transition matrices, difference matrix and their corresponding AACP. The AACP value of real data was then compared to the distribution of randomly shuffled trials. We observed that the AACP value of real data was beyond the 95th percentile of the distribution of shuffled data (Fig 6D, S8D), suggesting that transitions of the network from one region of the state space to another are plastic in nature.

### Intra-state Transitions are More Plastic than Inter-state Transitions

We further examined which transitions on the state space are significantly altered across sleep trials. We computed AACP specifically for transition from REM/non-REM/intermediate sleep state to REM/non-REM/intermediate state. We found that transitions occurring from REM-to-REM sleep and non-REM-to-non-REM sleep (intra-state transitions) are more vulnerable to plasticity after exploration as compared to inter-state transitions (such as non-REM to REM, REM-to-intermediate etc.) (Fig 6E, F). These changes to intra-state transitions were observed to be beyond randomness (Fig S8 E, F) suggesting non-uniform nature of plasticity in state transitions after exploration. We next investigated how fast the network flows on the state space and assessed whether the speed is uniform, or it exhibits specific region-dependent characteristics.

### Stabilization on State Space during Transition to REM

Coverage speed determines how fast the network sweeps the area of the state space. To calculate the coverage speed, we binned the state space as shown in Fig 7A, B and computed the number of bins the network covers in each time window (see Methods). Thus, the resultant time series provide information about how network’s flow varies over time. When speed is closer to its minimum value (i.e. 1 bin/s), the network stabilizes to a set of states which are closer to each other, whereas when speed approaches its theoretical maximum value, the network covers wider range of area, thus more potential fluctuations in the power of oscillations. We observed a reduction in median coverage speed during awake as compared to sleep trials (Fig 7C, S9B). During awake trials, due to subspace occupancy and behavioral constraints on state space, the network operates in constrained space as compared to sleep. Thus, the network is more likely to fall on states closer to each other, reducing its coverage speed as compared to sleep trials. We further explored the coverage speed during sleep, wondering how coverage speed varies during network transitions to REM sleep. We addressed this issue by taking a closer look at network trajectories and coverage speed when the network makes the transition (Fig 7D). We observed a sharp dip in coverage speed associated with state transition to REM sleep (Fig 7F, S9A). This further suggests that during the transition to REM sleep, the network stabilizes on state space.

**Fig 7.**
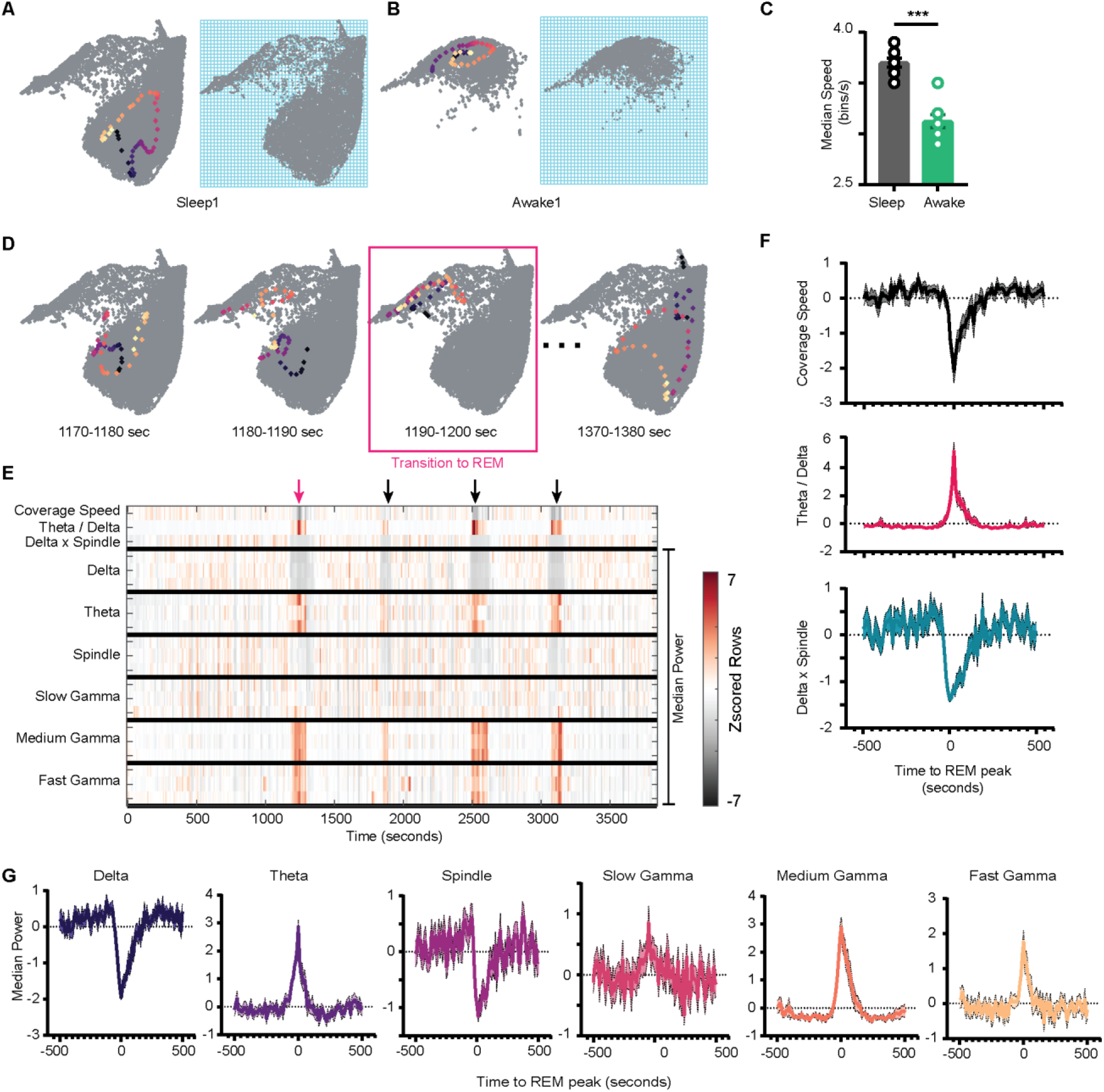
Network Stabilization during Transition to REM. A) Left: representative trajectory in 10 second timeframe on state space during sleep trial (trajectory starts from blue to yellow). Right: Binned state space used to compute the speed. B) Same as A, but for awake trial. C) Median speed for sleep and awake trials (n = 8 awake and sleep trials from 4 animals) (Sleep: 3.7 ± 0.04 v/s Awake: 3.12 ± 0.06 bins/s), p = 0.0003, Mann-Whitney test. D) Representative trajectories in 10 seconds timeframe before (panel 1 and 2) network transitions to REM state (3rd panel, red box), and when network exits REM (4th panel). E) Median power v/s time for entire sleep trial with 18 oscillations, speed of coverage, theta/delta ratio and delta x spindle product. Arrows indicate network transition to REM. Red arrow corresponds to example shown in D. Each row is independently z-scored. 3 rows for given oscillation corresponds to layers (pyramidal, radiatum and SLM respectively). Note the changes in median power of oscillations as network transition to REM. F) Summary plot for 19 REM bouts collected across 8 sleep trials from 4 mice. Top: Speed of Coverage Center: Theta/Delta Ratio Bottom: Delta x Spindle power product showing dip in coverage speed as network transitions to REM. G) Average power for 6 oscillations from pyramidal layer for 19 REM bouts as network transitions to REM. All are statistically significant at T = 0 sec when compared with T = −500 sec except slow gamma, p < 0.05, Mann-Whitney test. See also Figure S9.

### Dichotomous Nature of Gamma during Transition to REM

We next investigated the characteristics of stabilization during transition to REM. We wondered how the power in frequency bands is re-organized during the transition. We investigated this by plotting a spectrogram of median power for all 18 frequency bands during the entire sleep trial (Fig 7E). This allowed to visualize how speed of coverage varies with median power in all oscillatory bands. By definition of REM, we observed an increase in theta and a reduction in delta power during the transition. However, this transition was also associated with a significant reduction of power in spindle frequency band and with an increase of medium and fast gamma power while slow gamma remained unchanged (Fig 7G).

In the following section, we explored two applications of this analytical framework. We first use the network state space as a canvas (or context) to map the firing (activation signatures) of cells from CA1 pyramidal layer.

### Distinct Sleep State Signatures for Cells with Different Firing Rates and Sparsity during Awake Exploration

We visualized the network modulation of CA1 cells by overlaying cell’s firing rate on network state space. These plots represent how each cell activates as a function of network states (Fig 8B, S10B) and allowed to visualize how distinct oscillations simultaneously regulate the firing of individual cells during wakefulness and sleep. Each cell has its characteristic firing pattern on the state space (activation signature), as it does in the arena during exploration (firing rate-map) (Fig 8A, S10A). We hypothesized that distinct activation signatures on state space might corresponds to distinct firing rate-maps during exploration. To investigate this, we projected sleep signatures of all recorded cells for a given animal on a 2D plane (see Methods). We refer to this projected space as *signature space* (Fig 8C) as each point on signature space represents a cell’s activation signature during sleep trials. We then overlaid mean firing rate (Fig 8D, S11A) and mean sparsity (Fig 8E, S11B) of cells, as computed during awake trials, on signature space. We observed distinction of sleep signatures for low firing cells in the arena as compared to the high firing ones (Fig 8F). Specifically, we observed that sparse firing cells during exploration exhibits preferential firing to non-REM states (from Fig 8F and 4A,B) and are therefore simultaneously modulated by high power in delta, theta, spindle oscillations as well as moderate power in slow and fast gamma oscillations (from Fig 3A). However, the high firing cells during exploration are differently orchestrated by simultaneously occurring sleep oscillations as evident in Fig 8F. Altogether, this suggests that distinct functional cells in hippocampal CA1 are tuned to distinct set of network states and have distinct activation signatures during sleep trials. The network may employ such switches (distinct network states), to precisely manipulate the firing of different cells allowing them to play state-dependent functional roles. Studying cell’s firing patterns on network state space (context of the network) may provide new insights on how cells are functionally embedded in a larger network.

**Fig 8.**
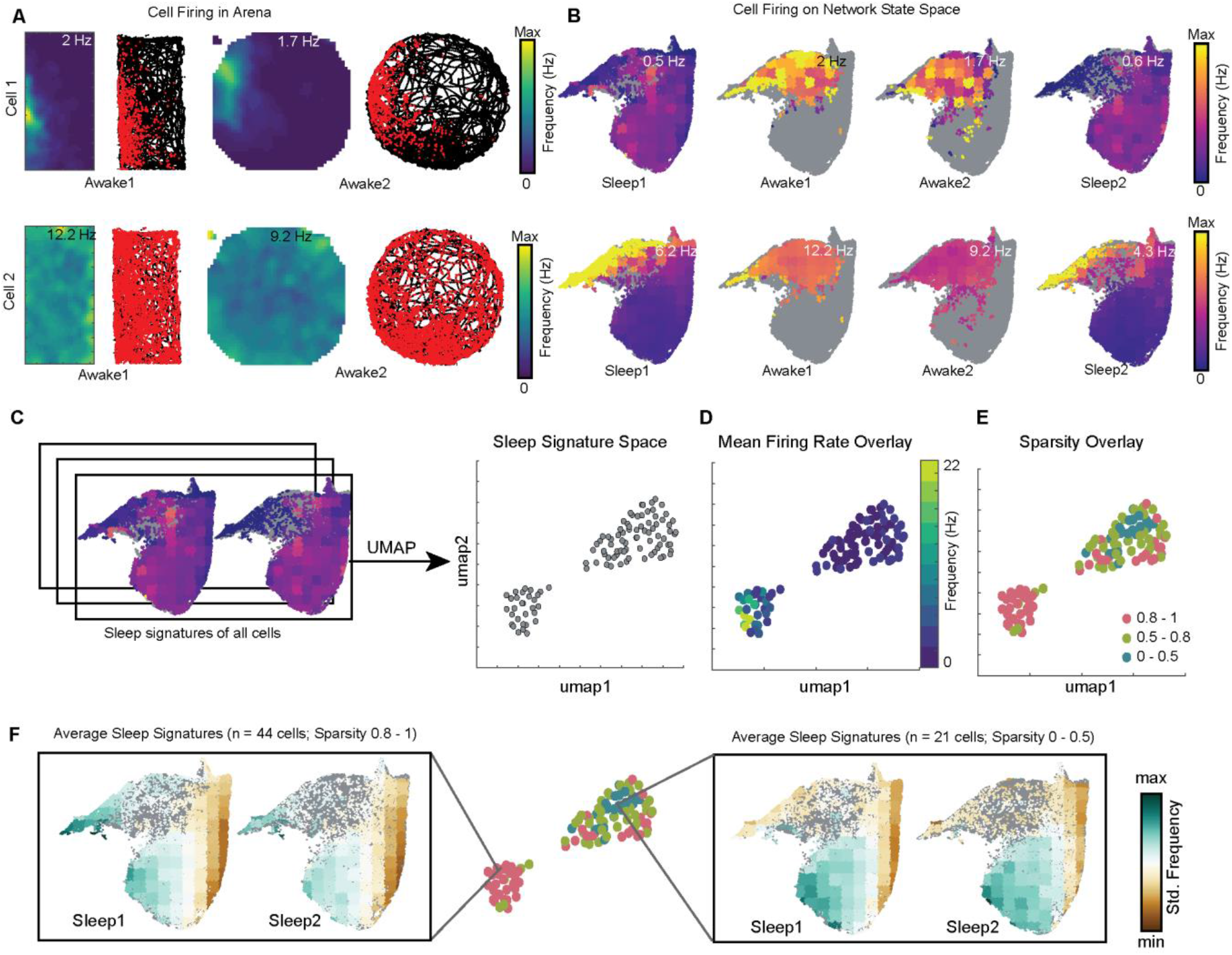
Distinct Sleep State Signatures for Cells with Different Firing Rates and Sparsity. A) Two representative cells and their firing during awake exploration trials (rectangle and circular arena). 1st and 3rd panel in both rows: Firing rate map for two trials. 2nd and 4th panel: animal’s trajectory during exploration (black) and overlaid with spikes (red). B) Firing of cells across 4 trials overlaid on network state space. Unvisited states are colored gray. Note distinct cells firing patterns in arenas corresponds to distinct cell firing patterns on network state space. C) Sleep signatures on state space are used to create signature space with 106 cells from a given animal. Each point represents firing signature of cell on state space across two sleep trials D, E) Mean firing rate and mean sparsity as observed during two awake trials, overlaid on sleep signature space. Note: Most high firing cells on the opposite end of the spectrum from low firing cells suggesting distinct sleep signature for cells having distinct firing rates and sparsity during awake exploration. F) Average sleep signatures for two set of cells having low (0 - 0.5) and high sparsity (0.8-1) during awake exploration. See also Figure S10 and S11.

### Altered Organization of Oscillations in the NLG3 knock-out Mouse, an Animal Model of Autism

We next applied the network state space approach to study the organization of network oscillations in the hippocampus of the NLG3 knock-out (KO) mouse, an animal model of autism spectrum disorders (ASD). NLGs are postsynaptic adhesion molecules which bind to their presynaptic partners neurexins to functionally couple the postsynaptic densities with the transmitter release machinery, thus contributing to synapses organization and stabilization (Südhof, 2008). Mice carrying the human mutations/deletions of the *Nlg3* gene, are useful animal models to study the mechanisms of ASD, since they recapitulate most of behavioral deficits found in autistic children. Previous studies from NLG3 KO mice have highlighted alterations in synaptic signaling and network oscillations, associated with social deficits (Földy et al., 2013; Modi et al., 2019; Tabuchi et al., 2007) Here, LFPs were used to extract network oscillations occurring at different frequencies from the CA1 region of awake head restrained NLG3 KO mice and wild type controls (WT) (Fig S12A).

We first constructed the composite network state space of hippocampal oscillations by pooling network states from all trials and all animals (Fig 12C, see methods, n = 7 trials from 4 WT and 5 trials from 3 KO mice). Each trial consisted of 3 minutes of LFP recordings while the animal was awake and at rest on a platform in a head-fixed condition (Fig 12A, B). Assessing the network state during rest allowed us to control for any genotype specific disparity in locomotion (Jaramillo et al., 2014; Modi et al., 2019) and its associated hippocampal oscillations. By overlaying trial specific states on the combined network state space (Fig 12C), we observed distinct occupation of state space by WT mice as compared to NLG3 KO mice. Overlay maps of power of different oscillations on the state space suggests that mice lacking NLG3 occupy state space associated with lower power of oscillations (Fig 12D, top). Comparing power of individual bands, reveals significant reduction in mean power of delta, theta, spindle, medium and fast gamma frequency bands. Only slow gamma power was not significant. We next computed correlation among various oscillatory processes (Fig S13 A, B) and observed no differences across genotypes. Lastly, we computed the speed of coverage and observed no difference across genotypes. Altogether, this suggests altered co-occurrence patterns along with intact coupling and transition properties of hippocampal oscillations in NLG3 KO mice (Fig S13C).

## DISCUSSION

In this study, we introduced an analytical framework for investigating the organization of hippocampal oscillations. This framework is established by constructing a state space of multiple network variables: in our case, the power of 18 hippocampal frequency bands from 3 layers of the CA1 region. We applied this framework to characterize various static and dynamic properties of the network state space during wakefulness and sleep and demonstrated multiple applications of the framework.

This multi-variate analytical approach provides several advantages over classical bivariate methods which aims to study only one oscillatory process at a time. By constructing the state space of power in various frequency bands from distinct layers of a brain region, our analysis allows to characterize the network landscape and, how various oscillatory processes co-vary as a function of the overall state of the network. Additionally, we demonstrated that the state space can be used as a canvas to map the activity of individual cells, thus allowing to study of how various oscillations can simultaneously influence the firing of neuronal ensembles. Although the scope of this study was limited to demonstrate the state space approach for power of network oscillations, the general idea can be extended to create a state space with additional attributes of oscillations such as instantaneous frequency, phase, etc. The resulting state space might provide additional insights about the organization of network oscillations. For instance, characterization of the state space constructed with instantaneous phase of the oscillations may provide novel information on how cell spiking is phase-locked to multiple oscillations simultaneously and how those phase-locking properties vary with time and behavioral state of an animal.

We first used the network state space to visualize the restricted nature of access to the state space during wakefulness as compared to sleep. We then characterized that restriction by overlaying the power of individual oscillations on the state space. This helped us to visualize how oscillations are intrinsically organized and co-vary simultaneously on the state space during wakefulness and sleep. As compared to sleep, in our experimental settings, during awake trials, the delta power is low, and the network is more dominated by theta and gamma supporting the ongoing navigational needs (Buzsáki and Moser, 2013; Colgin and Moser, 2010). Interestingly, it was recently reported (Furtunato et al., 2020) a specific increase in delta power accompanied by decrease in theta and gamma power in rats trained to execute 35 successive short-term treadmill runs at the same speed. These results suggest, as compared to our experiments, exploration of a larger subspace, at least in part related to changes in internal physiological states during performance of running exercise. We also applied the analysis to the datasets obtained from rat’s hippocampus and observed remarkable similarity across species (Fig S5), suggesting that the restrictions on network state space during wakefulness are conserved across species. Together, these observations indicate that, during wakefulness the hippocampal network occupies a fraction of subspace, independently of task and behavioral needs.

We next identified REM and non-REM states on the network state space and observed increased non-REM density during post exploration sleep trials. This finding complements a previous report (Eschenko et al., 2008) of sustained increased in sharp wave ripples activity during slow wave sleep after learning. In our case, the animal explores two novel environments during exploration trials, subsequently in the post-exploration sleep, the network spends more time on the non-REM states, possibly supporting the occurrence of sharp wave ripples and memory consolidation. Along with increase in the occurrence of high delta and high spindle power states (definition of non-REM), we also observe simultaneous increase in occurrence of other network oscillations such as theta, slow and fast gamma in post-exploration sleep trial (sleep2). The slow and fast gamma oscillations are linked to inputs from CA3 and medial entorhinal cortex, respectively (Colgin et al., 2009; Fernández-Ruiz et al., 2017), thus an increase in density of states characterized by medium and fast gamma might suggest an increased hippocampal-entorhinal and CA1-CA3 communication during non-REM sleep after exploration or learning. This modulation of active oscillatory states across sleep stages is also consistent with observed reorganization of cellular excitation between REM and NREM sleep (Grosmark et al. 2012, Miyawaki et al. 2016, 2019) which might mirror the differential inputs impinging on the network. Nevertheless, the precise mechanisms and functional role(s) of such increased inter-areal communication after exploration or learning remains to be determined.

Further, we characterized the coupling of various frequency bands during awake and sleep trials and across distinct network states. We reported state dependent coupling of distinct hippocampal oscillations suggesting that their sources change their functional link during distinct states on state space. For instance, during REM sleep, we observed spindle-frequency range oscillations coupled with slow gamma power and medium gamma coupled with fast gamma. We also observed delta decoupled from medium and fast gamma. These observations are in agreement with a general understanding of REM sleep as corresponding to a state of low-coherence across the cortex and between the cortex and the hippocampus (Diekelmann and Born, 2010) while arguing for the existence of frequency-specific channels of communication. In fact, this configuration alters during other network states such as non-REM sleep (Fig C). Besides the modulation of power of individual oscillations, the observed variations in coupling across frequency bands amplitudes might indicate a wider spectrum of combinations available to the network to dynamically regulate the activity of hippocampal population. Furthermore, such coupling states could mirror a similarly complex processing of specific inputs from up or downstream regions during distinct states of sleep and wakefulness (Colgin, 2015; Dvorak et al., 2021).

We next investigated the state transitions on the network state space. We quantified trajectories on state space and observed that state transition probabilities vary between two sleep trials and two awake trials. During exploration trials, the flow of the hippocampal network state is strongly affected by ongoing spatial navigation. The incoming and outgoing state-space trajectories are in fact bounded by the details of the exploration activity statistics (such as speed). Changes in transition probabilities thus reflect the alterations in behavioral sequences exhibited by the animal during exploration. During sleep trials, the state of the hippocampal network flows without any interference from environmental inputs. The incoming and outgoing trajectories bounds are instead characteristics of specific sleep states. Thus, changes in transition probabilities across sleep trials represent modifications in the probable paths on the network state space. These changes might be due to learning and/or experience acquired during awake exploration trials. The experience of running on a track or in an arena may change the state of the circuit in many ways. To give three possible mechanisms, (i) by changing the neuromodulatory state due to arousal, physical exertion. (ii) by changing brain temperature, which is known to modify brain potentials (Moser et al., 1993). (iii) by imprinting a trace of the experience by synaptic plasticity. Our methods enable to capture theses changes in terms of the coordinated modifications in oscillatory phenomena. Further experiments will be needed to disentangle the possible causes of these changes and how these dynamical changes correlate with changes in information processing in the hippocampus

We then characterized the speed of coverage on the network state space and reported the stabilization of network during transitions to REM sleep. We also described how frequency bands are re-organized during the transition. We observed specific increase in power of medium and fast gamma, but not slow gamma, during the transition to REM. State-dependent cross-frequency coupling between theta and gamma was also reported during REM sleep in EEG recordings from the mouse parietal cortex (Scheffzük et al., 2011). These results suggest the presence of common organizing principles across cortical and sub-cortical areas. It is worth mentioning that the slow gamma activity, which is not affected by the transition to REM, originates not only in the CA3 region of the hippocampus but also in the CA2 (Alexander et al., 2018; Middleton and McHugh, 2016). CA2 shows distinct anatomical (Hitti and Siegelbaum, 2014; Kohara et al., 2014) and functional (Dudek et al., 2016) properties which may differently affect REM sleep-dependent information processing.

Further, we demonstrated two applications of this analytical method. Firstly, rather than using individual oscillations to study the firing properties of hippocampal neurons, we overlaid the firing of hippocampal neurons on state space and used those sleep signatures to distinguish cells in CA1. We reported a gradient in the sleep state space signatures of cells, differentiating cells with sparse activation in the arena, from more active ones. This suggest that the sparse firing cells in the hippocampus are modulated differently by network oscillations occurring simultaneously during sleep.

Secondly, this method was applied to study the organization of hippocampal network state space in mice lacking NLG3, animal models of autism, which exhibit social deficits reminiscent of those found in autistic children (Bariselli et al., 2018; Modi et al., 2019; Radyushkin et al., 2009). In line with data obtained from animal models of ASD (Hammer et al., 2015; Modi et al., 2019; Paterno et al., 2021) or from the EEG/MEG of autistic children (Cornew et al., 2012; Larrain-Valenzuela et al., 2017; Ortiz-Mantilla et al., 2019; Rojas and Wilson, 2014; Wang et al., 2013), we observed a significant reduction of delta, theta, spindle, medium and fast gamma activity in NLG3 KO mice. These data are similar to those reported in the CA2 region of the hippocampus of NLG3 KO anaesthetized animals (Modi et al., 2019), suggesting that alterations in network activity in CA2 may influence the downstream CA1 network activity (Hitti and Siegelbaum, 2014; Kohara et al., 2014). However, the reduction in slow gamma power detected in CA2 (Modi et al., 2019) was not associated to a similar decrease in CA1 reported here, suggesting slow gamma oscillations in CA1 can emerge from sources other than CA2, possibly CA3 (Colgin et al., 2009).

Oscillatory activities, and in particular the gamma ones depend on the synchronous firing of large neuronal ensembles of excitatory cells within and across brain regions, which are paced by GABAergic interneurons (Gonzalez-Burgos and Lewis, 2008). Therefore, the observed alterations in power may be due to a reduced GABA release from GABAergic interneurons such as those containing parvalbumin (PV+) and cholecystokinin (CCK+) known to contribute to generate theta and gamma rhythms (Klausberger et al., 2005; Tukker et al., 2007). Interestingly, in a previous study from NLG3 KO mice (Földy et al., 2013), a clear increase of GABA release from CCK+ GABAergic interneurons into CA1 principal cells was detected, due to the impairment of tonic endocannabinoid signaling. Additionally, an impairment of GABAergic currents was also reported in CA2 region of NLG3 KO mice (Modi et al., 2019), an upstream region to CA1 (Hitti and Siegelbaum, 2014; Kohara et al., 2014). Therefore, the reduced oscillatory activity reported here is probably mediated by the loss of GABA_A_-mediated neurotransmission in both CA1 and CA2 hippocampal regions. This would cause an enhancement of network excitability with consequent excitatory/inhibitory unbalance, critical for controlling spike rate and information processing.

Lastly, we highlight some other potential applications of the network state space framework: *(i) Assessment of Inter-areal communication.* The state space approach can be applied to study oscillations from multiple brain regions simultaneously and how one set of oscillations from one region influence the others. *(ii) Analysis of EEG datasets.* Various oscillations can be extracted from EEG recorded from multiple brain areas and used to assess how they are organized during various behavioral tasks during wakefulness (i.e. decision making, navigation, imagination, mental calculation) or during sleep. Additionally, this analytical framework can be applied to study alterations in the organization of brain-wide network oscillations as signatures or biomarkers in various neurodevelopmental and neurodegenerative disorders. *(iii) Network Assessment of Novel Therapeutics*. Newly developed drugs are required to be extensively tested for their harmful side effects. Using network state space method will allow assessing how multiple brain regions, and their oscillations are simultaneously affected under pharmacological treatment. Additionally, this information can be used to predict potential side effects and develop better therapeutics in drug development.

## DATA AND SOFTWARE AVAILABILITY

The codes used for analysis are available at: https://github.com/brijeshmodi12/network_state_space Datasets will be made available upon request to F.P.B

## ACKNOWLEDGEMENTS

This research was supported in part by grants from the European Union’s Horizon 2020 research and innovation program (MGATE, grant agreement no. 765549 to M.G^1^. and F.P.B; BrownianReactivation grant agreement no. 840704 to F.S), European Research Council (ERC) Advanced Grant “REPLAY-DMN” (grant agreement no. 833964) to F.P.B; from Telethon (GGP 16083) and Del Monte Foundation grants to E.C; from EMBO to B.M (STF No. 8464). The authors are grateful to Dr. Bryan Souza for his help with programming. Rafael Pedrosa and Dr. Ashley Kees for their inputs regarding experiments in head-restrained mice. Prof. Peter Scheiffele for providing NLG3 KO mice. Dr. Gyorgy Buzsaki for providing the datasets in rats via Crcns.org. Dr. Michele Giugliano, Dr. Maurizio Mattia, Dr. Silvia Marinelli, Prof. Hannah Monyer and Prof. Antonino Cattaneo for their valuable discussion and suggestions on the research.

## AUTHOR CONTRIBUTIONS

B.M conceptualized and implemented the analysis with inputs from F.P.B and F.S. F.P.B. and M.G^1^ designed the experiments in freely moving mice. M.G^1^ conducted the experiments in freely moving mice, preprocessed the data and performed histology. B.M, M.G^2^ and E.C conceptualized the experiments from head-restrained mice. B.M conducted the experiments and analyzed the data from head-restrained mice. B.M wrote the original draft of the manuscript. E.C., F.P.B and F.S reviewed and edited the manuscript with additional inputs from all the authors. F.P.B supervised the freely moving experiments and analysis. M.G^2^ and E.C. supervised the experiments and analysis in head-restrained mice. All authors participated in the interpretation of the data.

## DECLARATION OF INTERESTS

The authors declare no competing interests.

## METHODS

### Ethics Statement

In compliance with Dutch law and institutional regulations, all animal procedures concerning recordings from freely moving or sleeping mice were approved by the Central Commissie Dierproeven (CCD) and conducted in accordance with the Experiments on Animals Act (project number 2016–014 and protocol numbers 0029).

All experiments from head-restrained animals were performed in accordance with the Italian Animal Welfare legislation (D.L. 26/2014) that implemented the European Committee Council Directive (2010/63 EEC) and were approved by local veterinary authorities, the EBRI ethical committee, and the Italian Ministry of Health (565/PR18) All efforts were made to minimize animal suffering and to reduce the number of animals used

### Animals

#### Recordings from freely moving mice

4 male C57BL6/J mice (Charles River) were used in this study, all implanted with a Hybrid Drive. All animals received the implant between 12 and 16 weeks of age. After surgical implantation, mice were individually housed on a 12-h light-dark cycle and tested during the light period. Water and food were available ad libitum.

#### Recordings from head-restrained mice

Experiments were performed on offspring male derived from heterozygous mating after 10 backcrossing with C57BL6/J. Results were analyzed blindly before genotyping. Control experiments were performed on wild-type littermates and C57BL6/J (WT). Genotyping was carried out on tail biopsy DNA by PCR using a standard protocol. After surgical implantation, mice were individually housed on a 12-h light-dark cycle and tested during the light period. Water and food were available ad libitum.

### Recordings from freely moving rats

Datasets of extracellular recordings from right dorsal hippocampus of 3 Long Evans rats using silicon probes were used for validating the findings. Detailed information is published in Grosmark and Buzsáki, 2016.

### Surgical Procedures and Data Acquisition

#### Extracellular recordings from freely moving mice

The fabrication of the Hybrid Drives and the implantation surgeries were done as described earlier (Guardamagna et al., 2022). These experiments were performed in Nijmegen, Netherlands. From post-surgery day 3 onward, animals were brought to the recording room and electrophysiological signals were investigated during a rest session in the home cage. Each day tetrodes were individually lowered in 45/60 µm steps (1/4 of a screw turn) until common physiological markers for the hippocampus were discernible (SWR complexes during sleep or theta during locomotion). Silicon probe signals were used as additional depth markers. Electrophysiological data were recorded with an Open Ephys acquisition board (Siegle et al., 2017). Signals were referenced to ground, filtered between 1 and 7500 Hz, multiplexed, and digitized at 30 kHz on the headstages (RHD2132, Intan Technologies, USA). Digital signals were transmitted over two custom 12-wire cables (CZ 1187, Cooner Wire, USA) that were counter-balanced with a custom system of pulleys and weights. Waveform extraction and automatic clustering were performed using Dataman (https://github.com/wonkoderverstaendige/dataman) and Klustakwik (Rossant et al., 2016) respectively. Clustered units were verified manually using the MClust toolbox. Manual waveform curation was performed using the MClust software suite. During all experiments in freely moving animals, video data was recorded using a CMOS video camera (Flea3 FL3-U3-13S2C-CS, Point Grey Research, Canada; 30 Hz frame rate) mounted above the arena. Animal position data was extracted offline using a deep learning based software, Deeplabcut (Mathis et al., 2018)

#### Extracellular recordings from head fixed mice

Separate set of mice were used for awake head fixed recordings at EBRI, Rome, Italy. 7 male mice (4 WT and 3 NLG3 KO) aged 3-5 months were implanted with stainless steel head bar (Luigs & Neuman, Germany). Mice were individually housed after implantation and were allowed to recover for 1 week. Mice were progressively habituated to head-fixation on a horizontal platform, until they no longer attempt to escape (3-4 weeks). For craniotomy, they were anesthetized with i.p. injection of a mixture of tiletamine/zolazepam (Zoletil; 80 mg/kg) and xylazine (Rompun, 10 mg/kg), a day before the recording. A glass electrode (Hingelberg, Malsfeld, Germany) with the resistance of 1–2 MΩ, filled with standard Ringer’s solution containing the following (in mM): 135 NaCl, 5.4 KCl, 5 HEPES, 1.8 CaCl2, and 1 MgCl2 was lowered to target CA1 pyramidal layer using micromanipulator (Scientifica, UK). Location of electrode was assessed by visually inspecting LFP trace, depth information and was confirmed post-hoc using brain sectioning and immunohistochemistry. LFPs were acquired with Multiclamp 700B amplifier (Molecular Devices, USA) and digitized with an A/D converter (Digidata 1550, Molecular Devices, USA). Data were acquired at sampling rate of 10 kHz and were further down sampled to 1KHz using MATLAB for all the further analysis of state space. Multiple 3-minute trials were recorded from a mouse and artifacts free trials were considered for all further analysis.

### Behavioral Paradigm

#### In datasets recorded from mice

In the open field experiments mice were free to explore two arenas for about 20 minutes each. The arenas were either square (45 cm x 45 cm) or circle (45 cm diameter) or rectangle (50 cm x 25 cm). This was preceded and followed by a rest session in the animal’s home cage (Sleep1 and Sleep2) that typically lasted 60 minutes each.

#### In datasets recorded from rats

Each session consisted of a long (∼4 hour) PRE rest/sleep (sleep1) epoch home-cage recordings performed in a familiar room, followed by a Novel MAZE running epoch (∼45 minutes) in which the animals were transferred to a novel room, and water-rewarded to run on a novel maze. These mazes were either A) a wooden 1.6m linear form, B) a wooden 1m diameter circular platform or C) a 2m metal linear platform. Animals were rewarded either at both ends of the linear platform, or at a predetermined location on the circular platform. The animal was gently encouraged to run unidirectionally on the circular platform. After the MAZE epochs the animals were transferred back to their home-cage in the familiar room where a long (∼4 hour) POST rest/sleep (sleep2) was recorded. Detailed paradigm and recording conditions are described in Grosmark and Buzsáki, 2016.

### Histology

After the final recording day tetrodes were not moved. Animals were administered an overdose of pentobarbital (300∼mg/ml) before being transcardially perfused with 0.9% saline, followed by 4% paraformaldehyde solution. Brains were extracted and stored in 4% paraformaldehyde for 24 hours. Then, brains were transferred into 30% sucrose solution until sinking. Finally, brains were quickly frozen, cut into coronal sections with a cryostat (30 microns), mounted on glass slides and stained with cresyl violet. The location of the tetrode tips was confirmed from stained sections (Fig S1).

### Neural Data Analysis

#### Preprocessing of Raw Signals

Movement and other artifacts were removed from LFP signals by setting an amplitude threshold (approx. > 6 median absolute deviation). Threshold were adjusted by visual inspection of raw traces. Data points with amplitude greater than threshold were eliminated from analysis pipeline. Raw LFPs recorded at 30 KHz were first lowpass filtered (<1000 Hz) and then downsampled to 1KHz for all further analysis of state space. Oscillations were extracted in 6 frequency bands: delta (1-5 Hz), theta (6-10 Hz), spindle (10-20 Hz), slow gamma (20-45 Hz), medium gamma (60-90 Hz) and fast gamma (100-200) Hz from each of 3 layers of CA1 using *eegfilt* from EEGLAB, an open source toolbox in MATLAB (Delorme and Makeig, 2004). CSD were computed from LFP using methods described in Lasztóczi and Klausberger, 2014.

#### Construction of Network State Space

Filtered LFP or CSD bands were used to compute power. Power time series were binned in 200 milliseconds time bins to compute median power. They were further smoothed using Gaussian kernel with standard deviation of 3 bins. The resulting smoothened, binned, 18 power time series were given as input to UMAP to obtain 2D projection with parameters: *nearest neighbors: 25, minimum distance: 0.1, distance metric: Euclidean*.

#### Combined state space from head fixed mice

LFPs from all WT and NLG3 KO mice were normalized by scaling from 0 to 1. The scaled LFP were then filtered to obtain the following frequency bands: delta (1-5 Hz), theta (6-10 Hz), spindle (10-20 Hz), slow gamma (20-45 Hz), medium gamma (60-90 Hz) and fast gamma (100-200 Hz). The median band power was computed in bins of 200ms for all 6 frequency bands. The network states from all the animals were merged and were projected into 2D using UMAP with parameters as described before.

### Identification of sleep periods

3-Axis accelerometer data were used to identify periods of mobility and prolonged immobility during recordings in home cage (Fig S2). Periods with mobility were excluded from further analysis.

### Fraction of State Space Occupied

The projected state space was divided into 100 x 100 bins. The occupancy was computed by counting number of bins visited during each trial. The visited bin count was then divided by total number bins in state space to get the fraction of state space occupied during each trial.

### Surrogate Data for Occupancy

The 18 binned power time series were randomly shuffled 100 times to create surrogate state spaces. UMAP projections were obtained for surrogate data and occupancy was computed. The measure *sleep-awake* occupancy was used to compare surrogate data with the observed data.

### Power Overlay on State Space

The projected state space was divided into 20 x 20 bins. For each trial and for each oscillation, median power in each bin was computed. The range of each oscillation was determined between minimum value and 95^th^ percentile to prevent skewness of sequential color scale by extreme artifacts. Sequential colors were assigned to power values in the determined range to generate overlay plots Fig 3. Unvisited bins in a trial were assigned gray color.

### Identification of REM and Non-REM states

We adapted the methods from previous study for identification of REM and Non-REM sleep (Grosmark et al., 2012; Mizuseki et al., 2009, 2011). The projected state space was divided into 20 x 20 bins. For each bin on state space, we computed median theta / delta power ratio of all the network states for identification of REM sleep states. For reproducibility, all the bins, having median theta/delta ratio greater than 80^th^ percentile was classified as REM. For identification of non-REM states on state space, we computed median delta x spindle power product for all the network states in a bin. All the bins, having median delta x spindle product greater than 70^th^ percentile was classified as non-REM. These were further verified by visual inspection LFP traces. The remaining states were classified as intermediate sleep states without any further sub-classification. The classified states were then visually inspected and cross-checked using accelerometer data and raw LFP traces. Oscillations from pyramidal layer only were used for identification.

### Density Overlay on State Space

The projected state space was divided into 20 x 20 bins. For each trial, we computed number of network state visits in each bin and then divided the count by trial duration to get normalized states count also known as density. Color assignment to density is as described above in power overlay methods.

### Correlation Matrix of Frequency Bands

Correlation matrices for each trial were created using *corrcoef* function in MATLAB. Input to *corrcoef* were trial specific power of 18 oscillations binned in 200ms bins (trial specific states). *corrcoef* computes pairwise Pearson correlation for all 18 oscillations for a given trial. For state dependent coupling of frequency bands, the projected state space was divided into 3 x 3 bins. For each bin, a correlation matrix was generated using the corrcoef. The inputs to corrcoef were bin-specific power of all 18 the oscillations (namely, all the states in that bin). To generate correlation matrix space, we took the lower triangular values of correlation matrices of all bins in all trials and used this as an input to UMAP. The embedding was generated using cosine distance as it brings similar points (in this case, correlation matrices) together. Thus, allowing us to separate similar bins. Each bin was then assigned its status based on the following: 1) Awake bin — Bins collected from awake trials. 2) REM bins — Bins from sleep trials having more that 10% of its network states identified as REM as described above. 3) Non-REM bins — Bins from sleep trials having more than 10% of its network states identified as non-REM as described above. 4) Intermediate — the remaining bins from sleep trials.

### Trajectories on Network State Space

The projected state space was divided into 3 x 3 bins for sleep trials and 10 x 10 for awake trials. For all the network states in each bin, we show the lines (trajectories) connecting 5 states (1 second of time) preceding and proceeding the state. We label set of preceding states as incoming trajectories and proceeding states as outgoing trajectories.

### State Transition Matrix

Transition probability from say, bin A to bin B was computed as following: for every state in bin A, we collected outgoing trajectory states (5th state from the current state, 1 second in future). Transition probability from A to B was then computed by fraction of total trajectory states that ended in B. Transition probabilities were then represented in the form of matrix.

### Change in Transition Probabilities and Average Absolute Change

Difference matrices were computed by subtracting corresponding bin transition probabilities from two sleep or two awake trials. Mean of absolute values from difference matrices were used to generate summary plot for all animals.

### Surrogate Sleep and Awake Trajectory Trials

State transition occurring in both the sleep (Sleep1 and Sleep2) and awake (Awake1 and Awake2) trials were merged to create a pool of state transitions. The sleep or awake pool were then split randomly into two surrogate sleep or awake trials respectively. 1000 randomly generated surrogate sleep or awake trials were generated to compute the change in transition probabilities between two surrogate sleep or surrogate awake trials and compute its average absolute change.

### Speed of Coverage on State Space

The projected state space was divided into 50 x 50 bins. We counted the number of bins visited by the network trajectory in a time frame of 10 seconds. Thus, the speed is measured in units: bins/sec. Median values of coverage speed from all time frame of sleep and awake trails were used to plot summary.

### Speed of Coverage during Transition to REM

Entire sleep trial was divided into non-overlapping time frames of 10 seconds. We then computed median theta/delta ratio for all the states in each time frame. REM peaks were identified by detecting peaks in median theta/delta ratio. Median power on state space from each oscillation in each time frame was also computed to generate spectrogram. Summary plots were generated by averaging power in oscillations from pyramidal layer, speed of coverage, theta/delta ratio and delta x spindle product for all REM peaks.

### Cell Firing Overlay on State Space and in the Arena

Spike trains of each cell were binned into non-overlapping bins 200 ms (same bin size as applied during the construction of network state space). Thus, each network state gets a corresponding bin of spike train. For generating overlay maps, the projected state space was divided into 15 x 15 bins. For each bin on state space, we identified a set of network states in that bin and computed their mean firing rate from corresponding bins in spike train. Plots demonstrating firing rate map in the arena were generated using methods described in (Fyhn et al., 2007)

### Firing Properties Overlay on Cell’s Signature Space

We computed the cell’s firing rate on sleep state space (sleep signature) as above. We used these sleep signatures of all cells of a given animal, as an input to UMAP with cosine distance and 10 neighbors. The resulting projected space is known as sleep signature space. We computed mean firing rate and mean sparsity of each cell during awake trials and overlaid it on sleep signature space.

## SUPPLEMENTARY DATA

**Fig S1.**
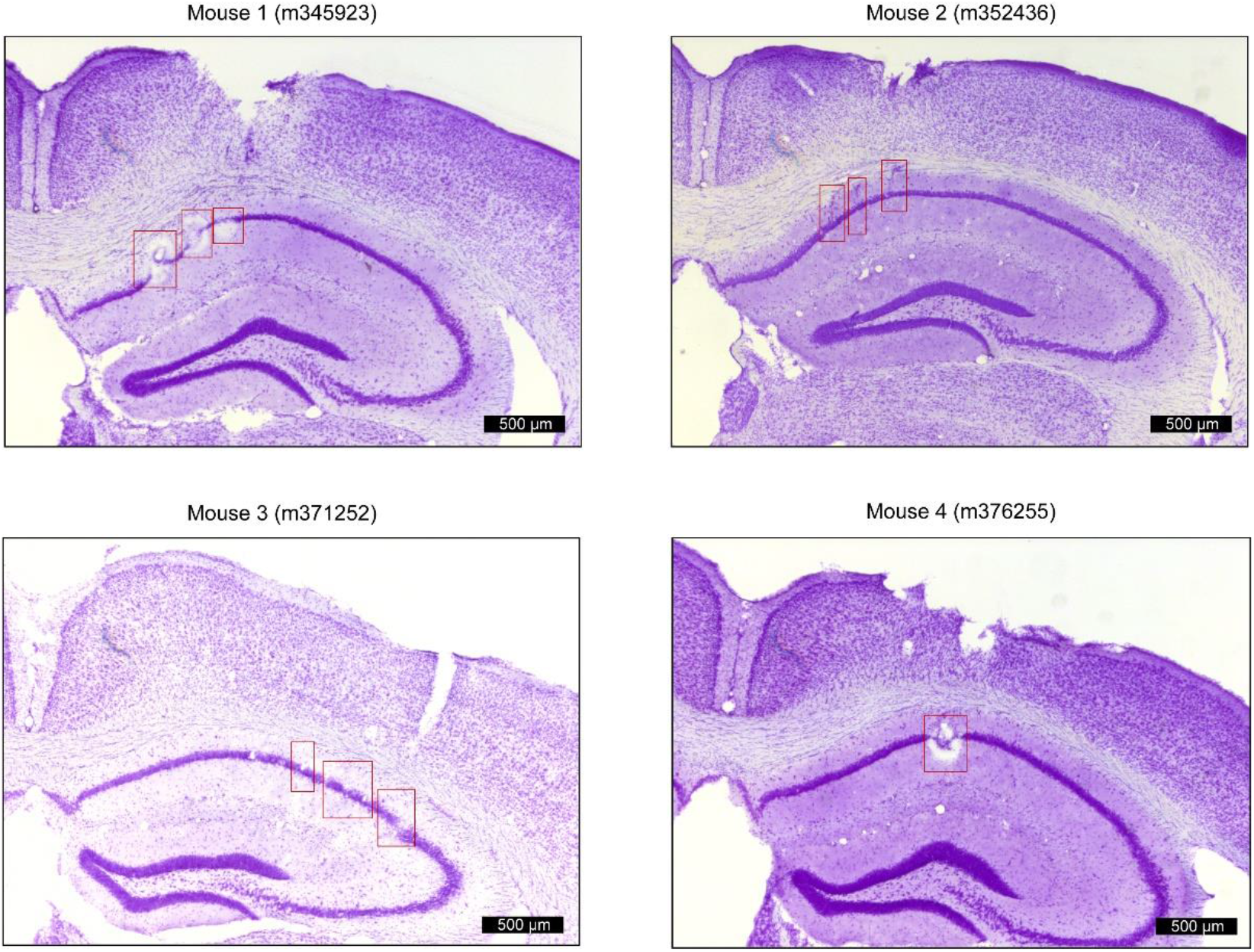
Positions of tetrodes targeting dorsal CA1 in freely moving mice. Four representative Nissl-stained coronal section showing recording locations from each animal (Mouse 1,2,3,4). Red squares indicate the estimated location (CA1 pyramidal layer) of tetrode tips after electrolytic lesions.

**Fig S2.**
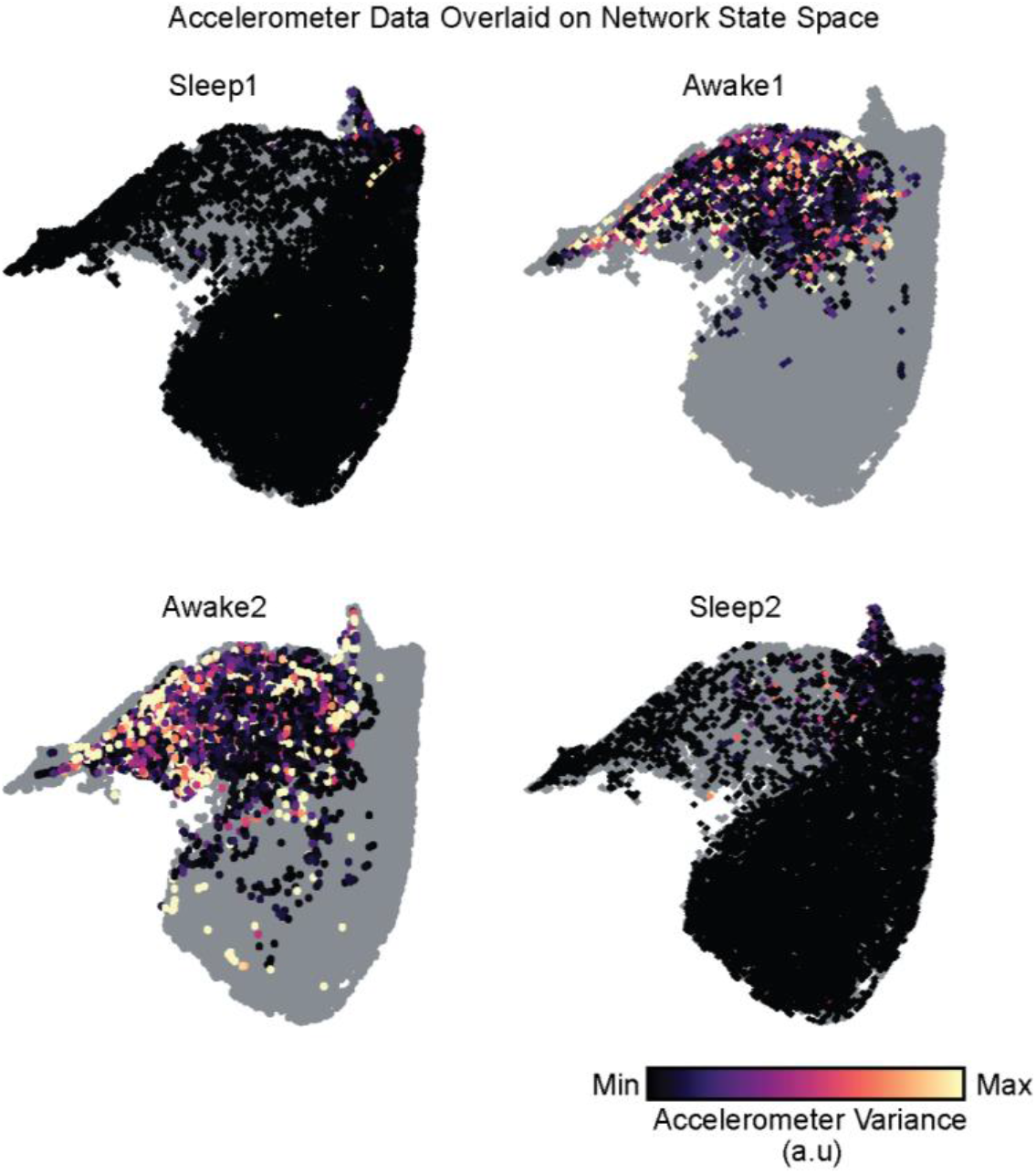
Accelerometer signal in arbitrary units overlaid on network state space, used for identification of sleep states during rest periods.

**Fig S3.**
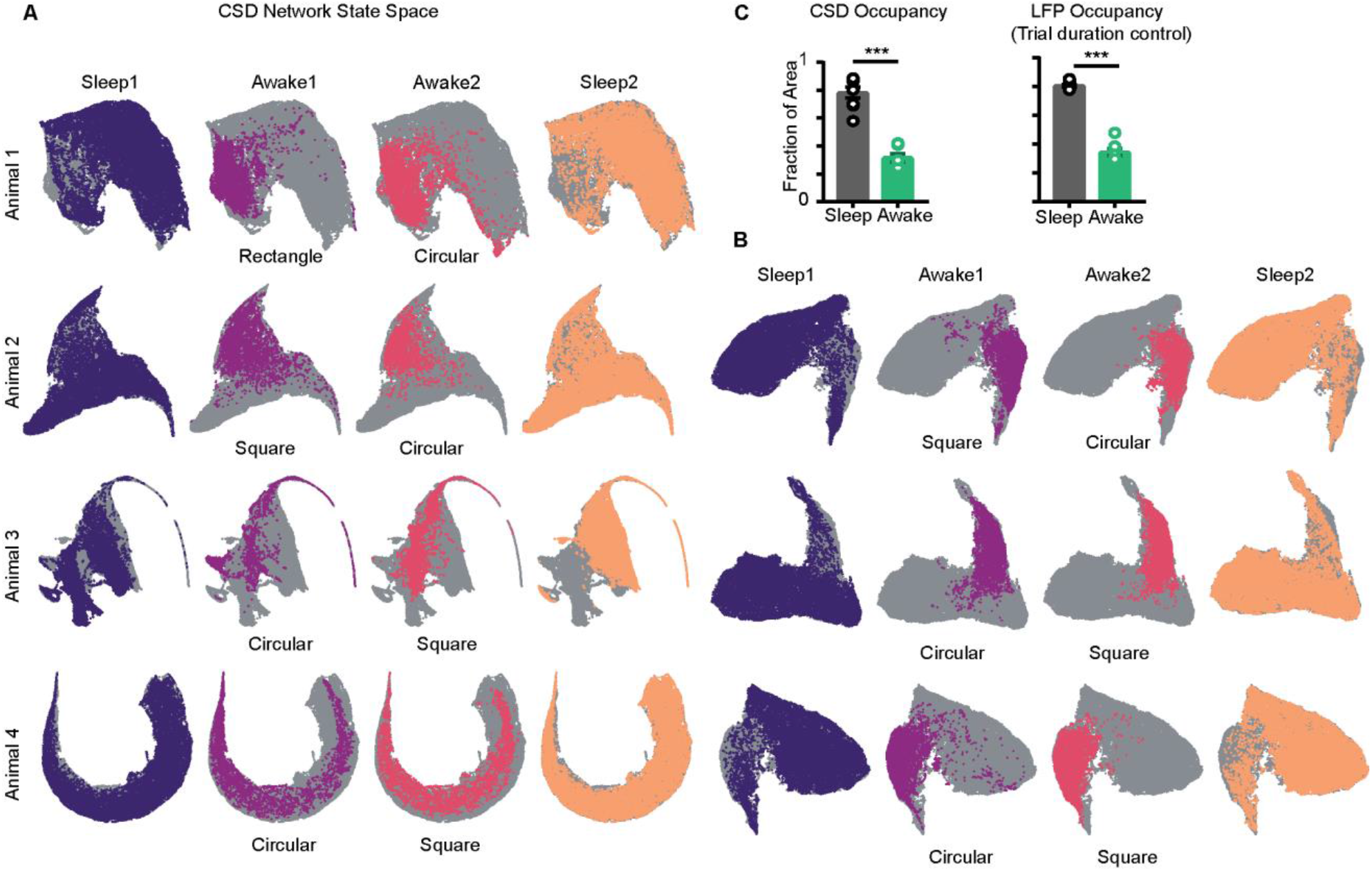
Trial specific states overlaid on CSD and LFP network state space. A) Network state space constructed using power in CSD signals, instead of LFP for 4 animals. Unvisited states are colored in gray. Trail specific states are colored. B) Network state space constructed using power in LFP signals for the remaining 3 mice. C) Left: Fraction of state space occupied on CSD state space Sleep v/s wake (0.78 ± 0.03 *versus* 0.3 ± 0.02) Right: Fraction of state space occupied on LFP state space obtained by combining equal number of states from sleep and awake trial, controlling for difference in trial duration (0.8 ± 0.01 *versus* 0.34 ± 0.02) (p = 0.0002, Mann-Whitney test).

**Fig S4.**
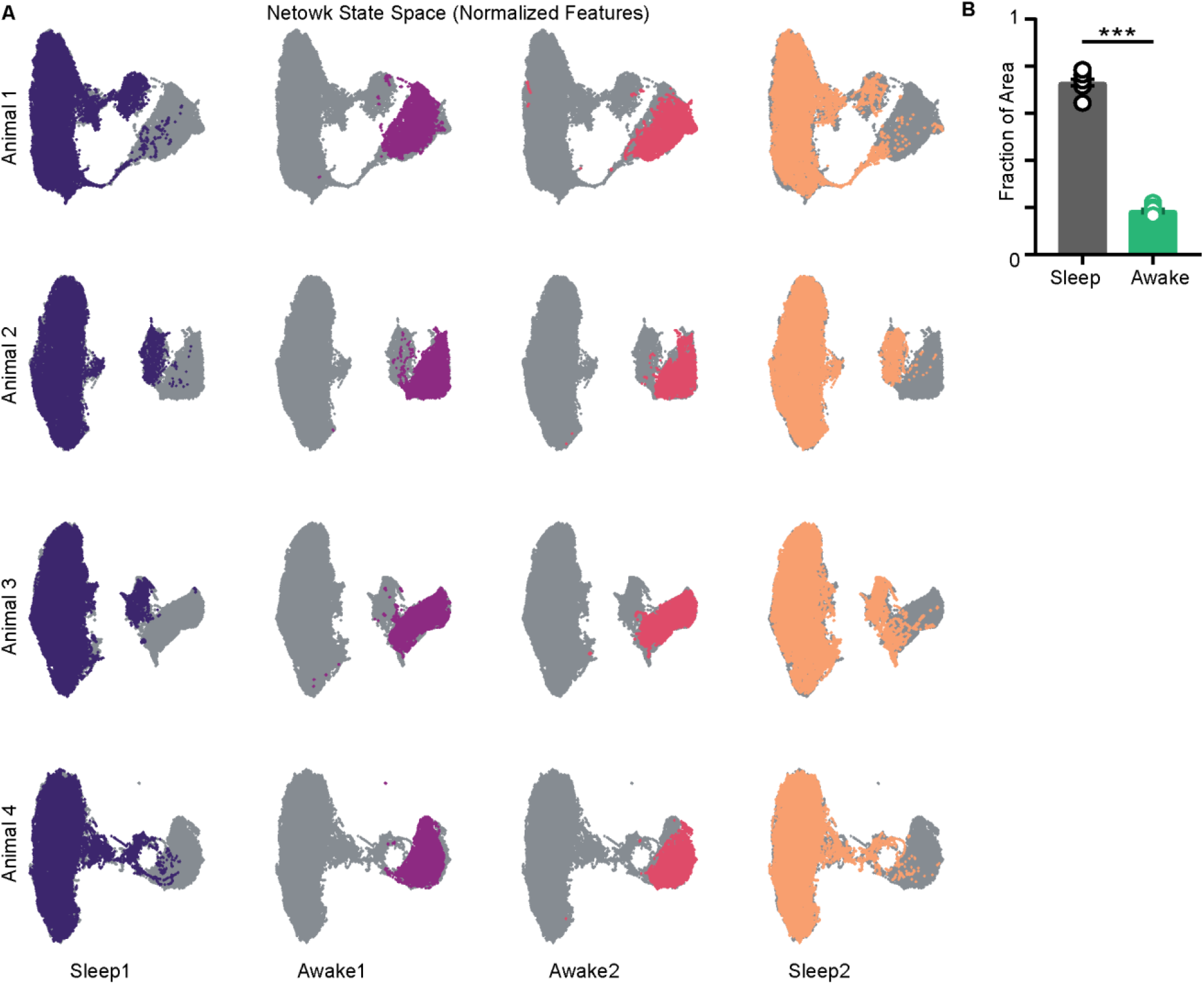
Network state space (LFP) computed with standardized features (z-scored) A) State space from 4 mice and trial specific states overlaid on it (colored) B) Fraction of area occupied on normalized state space Sleep v/s Awake (0.73 ± 0.01 *versus* 0.1 ± 0.007), p = 0.0002, Mann-Whitney test.

**Fig S5.**
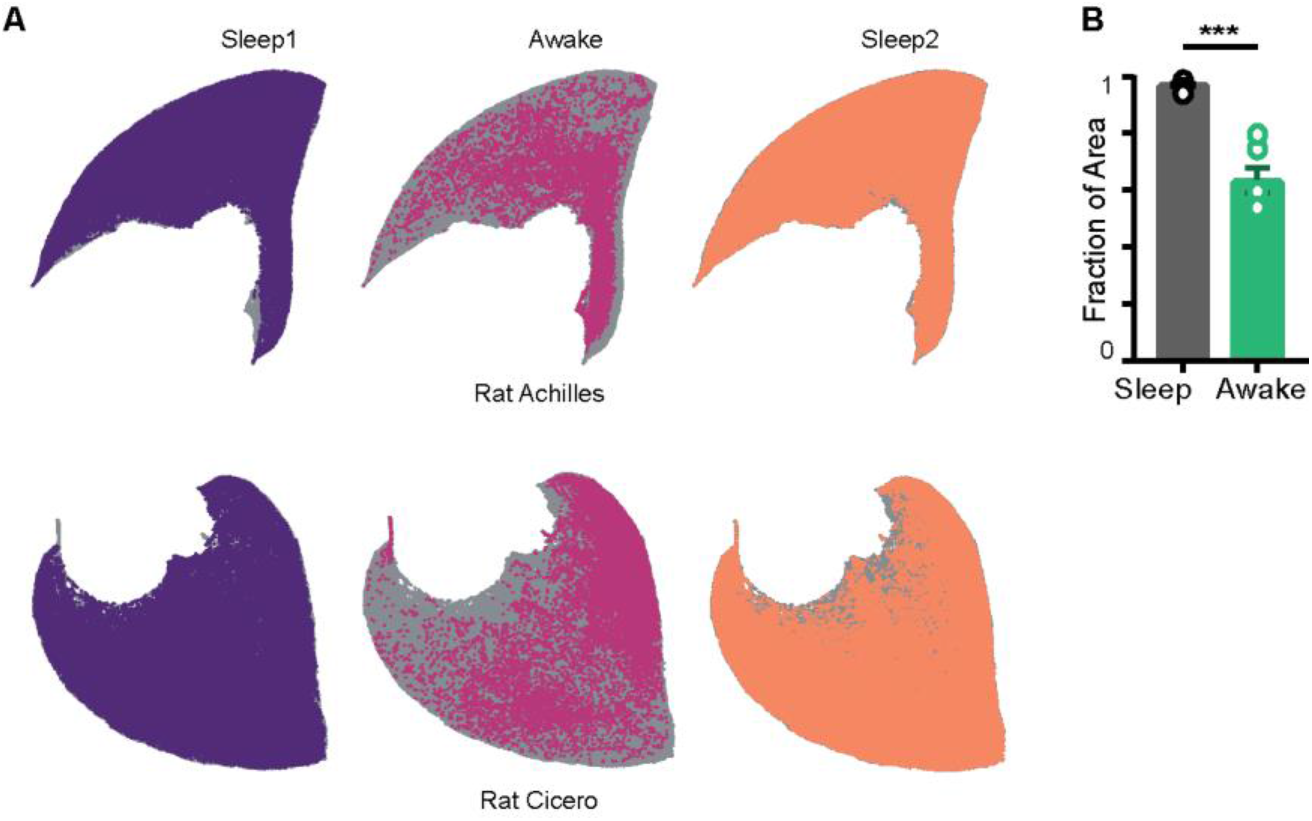
Restricted occupancy on the network state space in Rats A) Representative network state space from 2 rats(gray) and states occupied during sleep and awake trials. Datasets obtained from crcns.org courtesy of Dr. Gyorgy Buzsaki’s lab at NYU. B) Fraction of area computed from rat datasets. Sleep v/s awake (0.97 ± 0.003 *versus* 0.5 ± 0.04), p = 0.0001, Mann-Whitney test. (n = 12 sleep and 6 awake trials from 3 rats)

**Fig S6.**
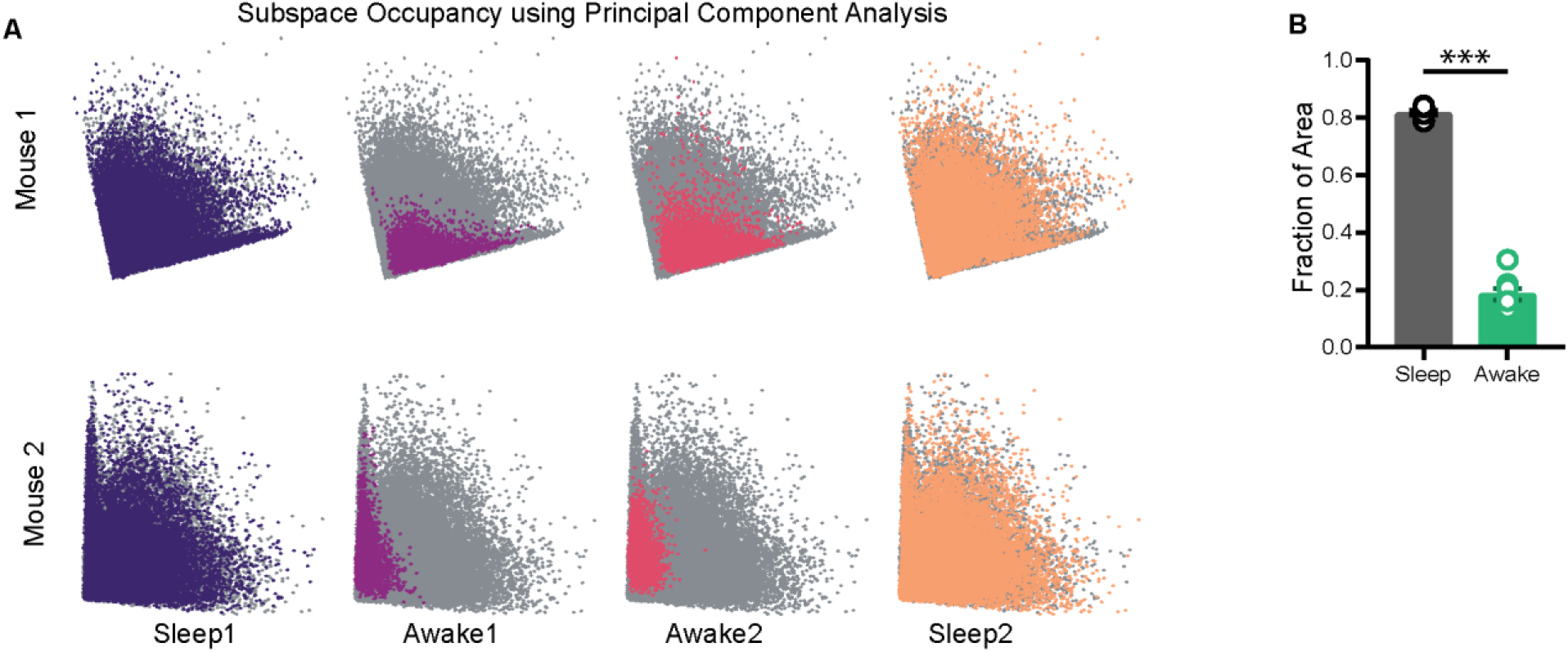
Subspace occupancy visualized using Principal Component Analysis. A) Representative network state space of 2 mice projected on 2D plane using principal component analysis (∼ 80 % variance explained). Note that the visualization of restrictions on the state space is independent of the dimensionality reduction method employed. B) Fraction of area occupied as computed on projections obtained using PCA. Sleep (0.81 ± 0.008) versus Awake (0.18 ± 0.02), p = 0.0002, Mann-Whitney test. (n = 8 sleep and 8 awake trials from 4 mice)

**Fig S7.**
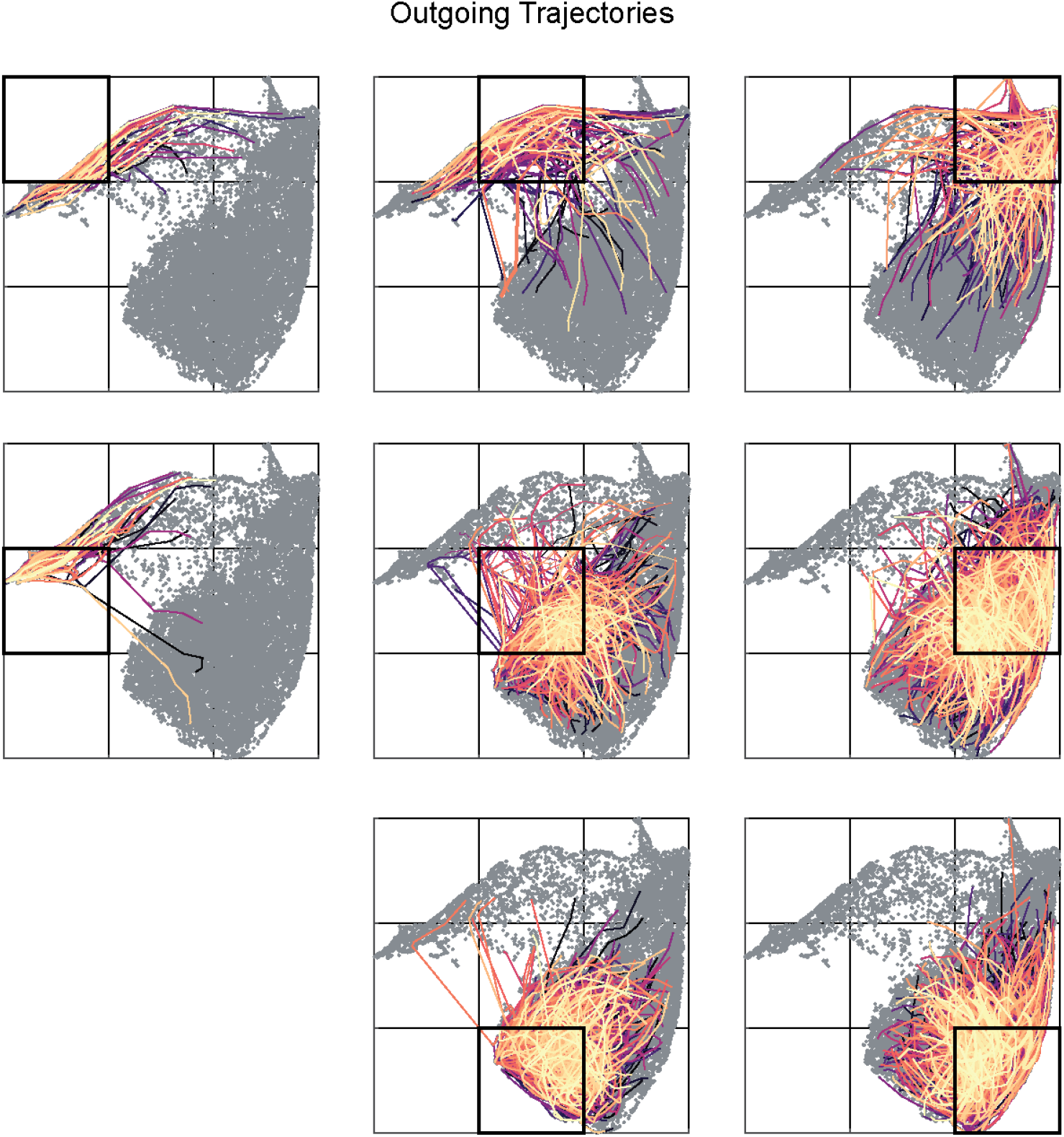
Outgoing trajectories (1 second in future) for all bin of a representative sleep trial. Distinct colors correspond to trajectories originating from distinct states in a given bin (highlighted in black)

**Fig S8.**
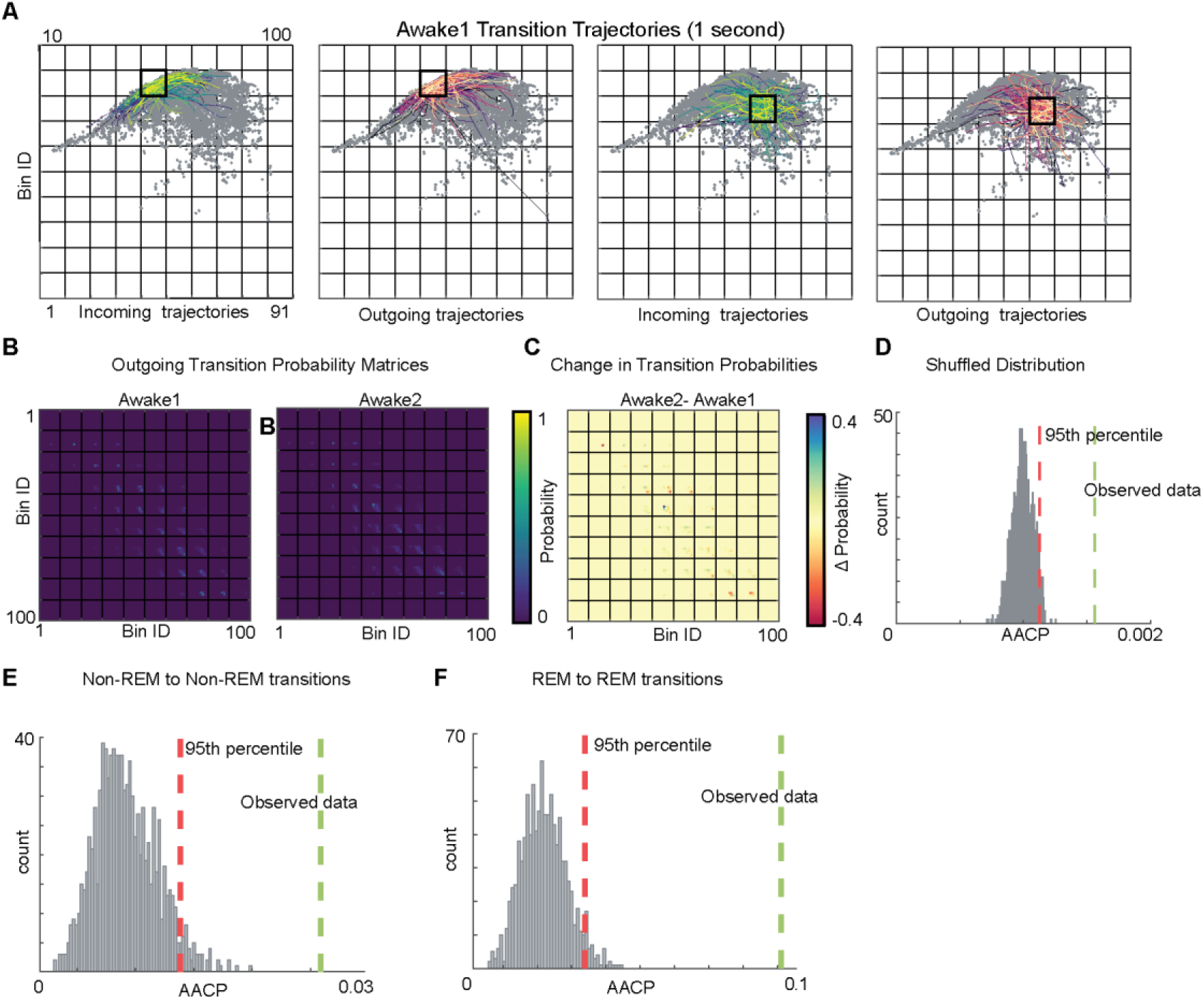
Altered Transition Probabilities across Awake Trails A) Incoming and outgoing trajectories for two representative bins on state space during an awake trial. Each trajectory spans 1 second in time before (incoming) and after (outgoing) the occurrence of given state in a given bin. B) State transition matrices for two awake trials. C) Change in transition probabilities across two awake trials obtained by subtracting two transition matrices (sleep2 - sleep1). D) Comparison of observed data’s average absolute change in probability with shuffled data. AACP for Real data = 0.0016, 95th percentile of shuffled data = 0.0011. E and F) Comparison of observed data’s AACP computed for REM-to-REM and Non-REM to Non-REM transitions across sleep trials with shuffled data.

**Fig S9.**
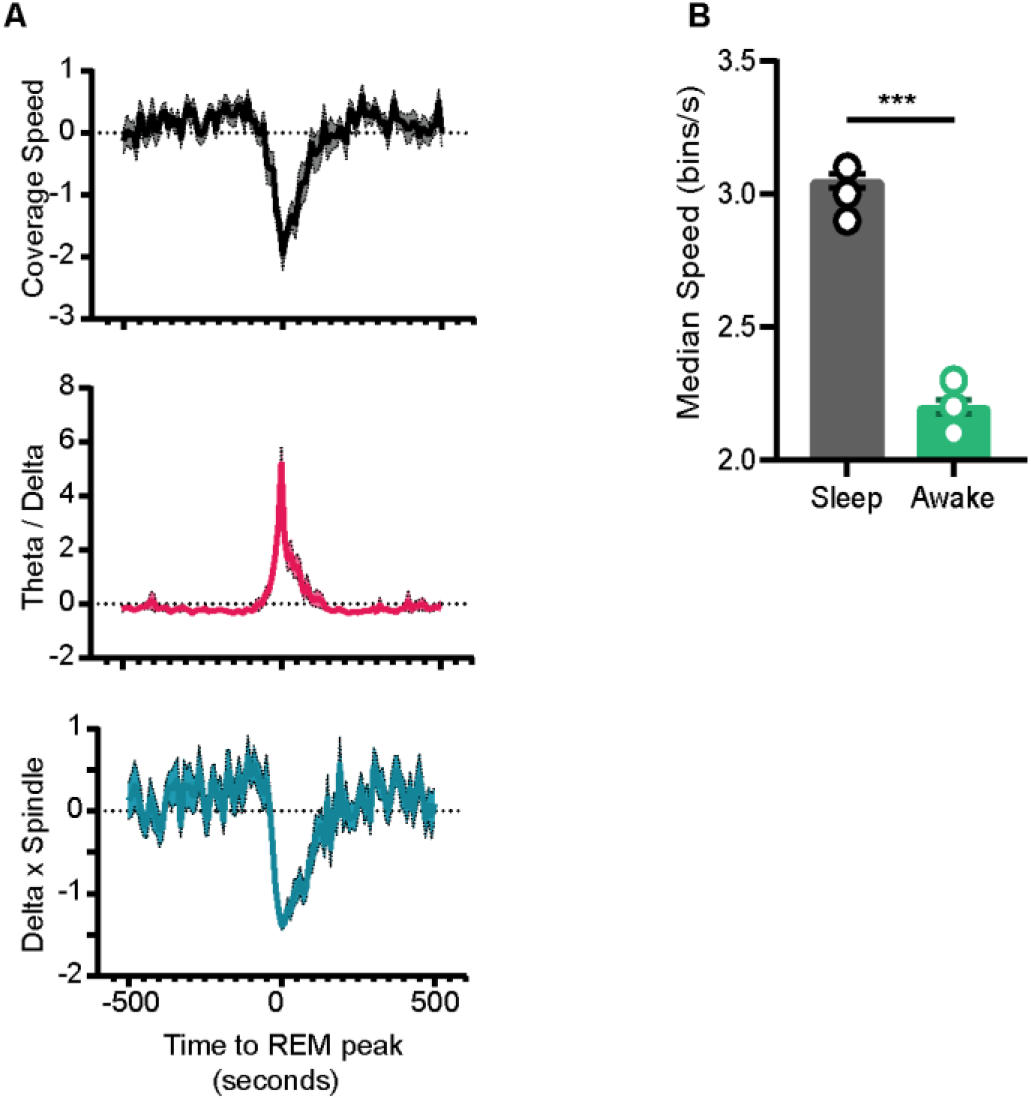
Speed of Coverage on Normalized State Space A) Top, Speed of coverage on normalized state space during transition to REM. Center and Bottom, Theta/Delta power ratio and Delta x Spindle power product (same as Fig 7F) B) Median speed of coverage computed on normalized state space for sleep and awake trials (Sleep v/s Awake 3 ± 0.02 *versus* 2.2 ± 0.02) bins/sec. p = 0.0002, Mann-Whitney test

**Fig S10.**
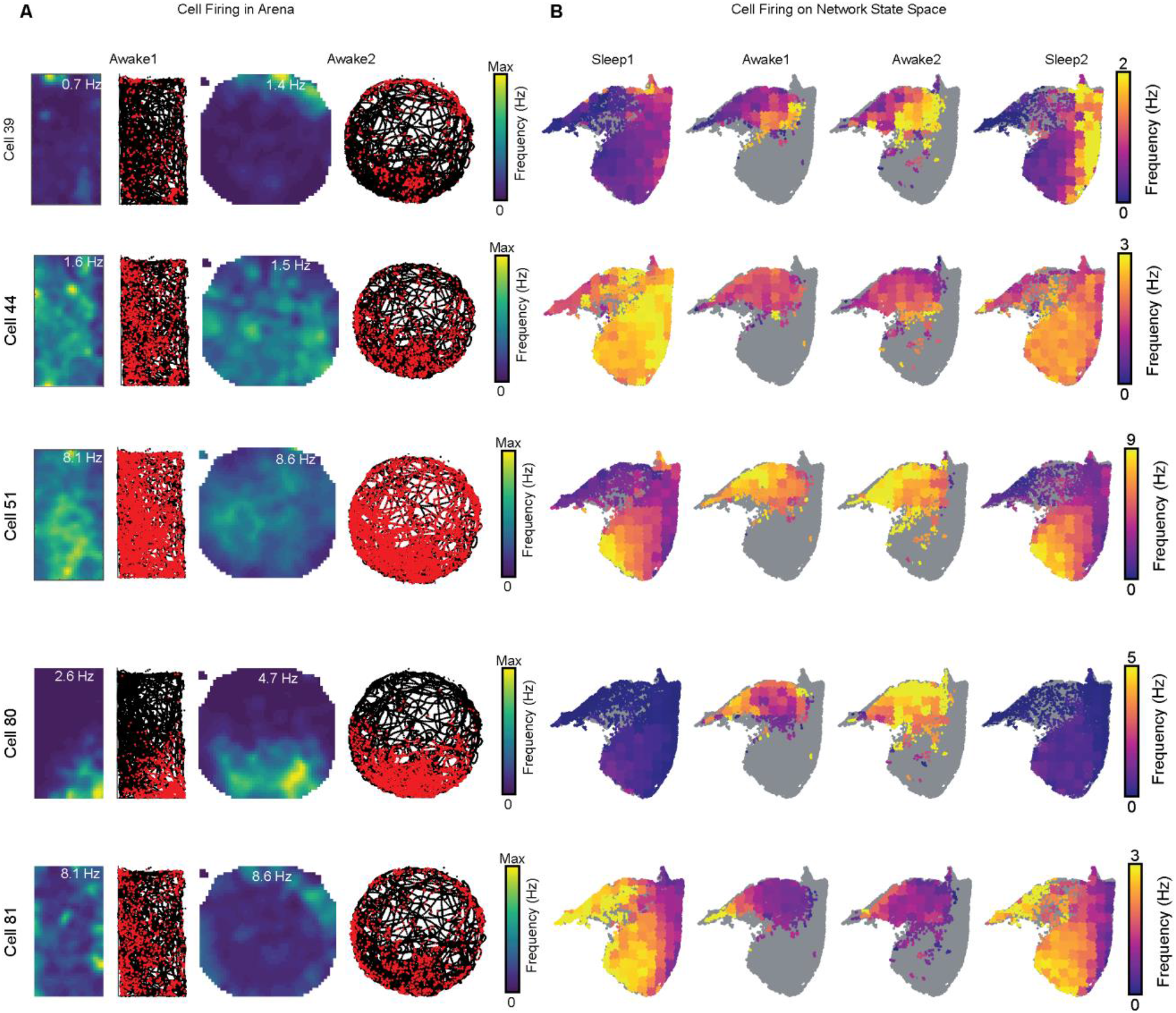
Cell Firing Overlaid on Arenas during Exploration and on the Network State Space A) Representative cells with rate maps, spikes (red) and trajectory(black) on two arenas during awake exploration. B) Corresponding cell firing rate overlaid on the network state space during 4 trials.

**Figure S11.**
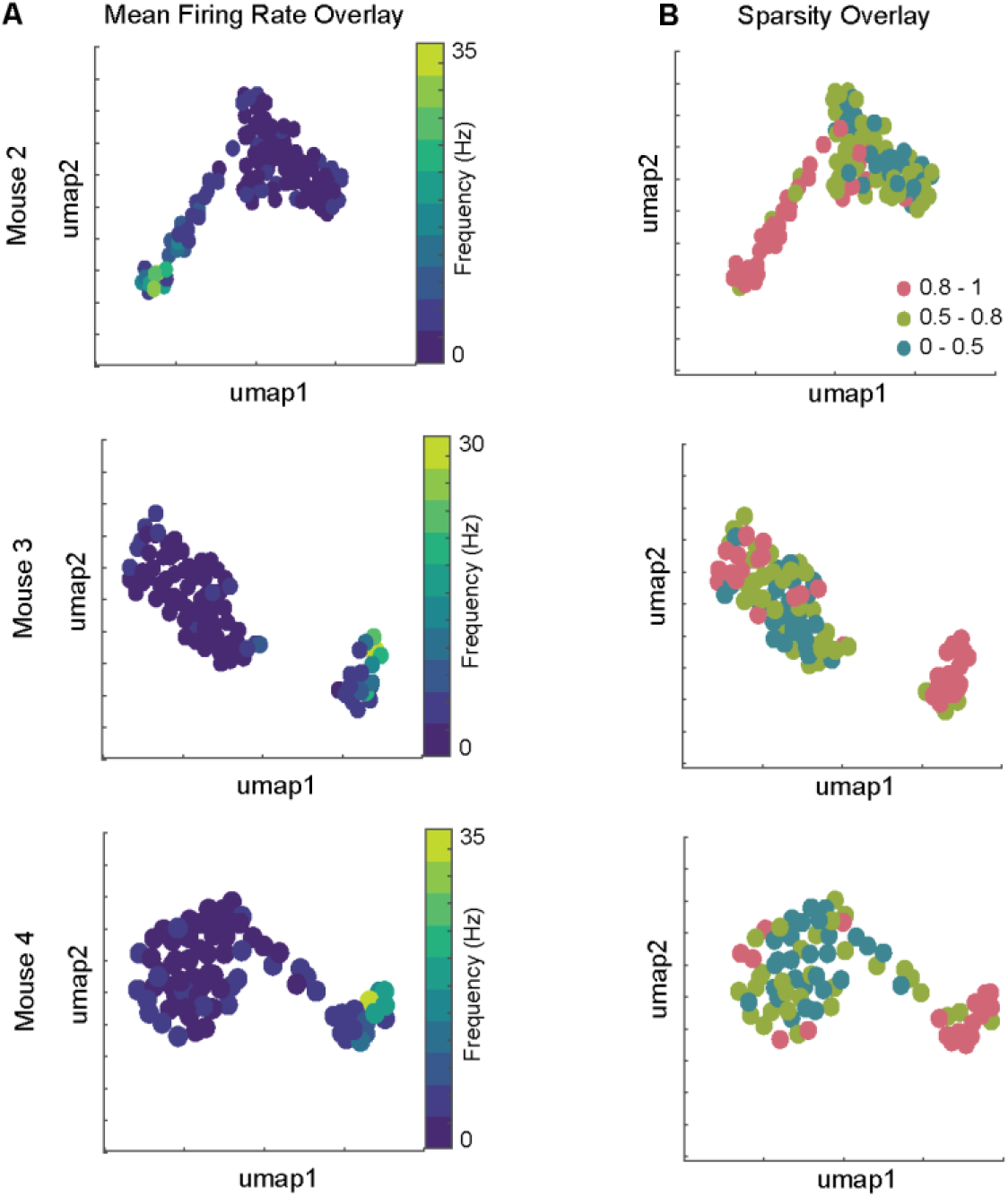
Distinct Sleep Signature corresponds to Cells with Distinct Firing Patterns in Arena. A and B) Mean firing rate and mean sparsity as observed during two awake trials, overlaid on projected sleep signature space. Total 115, 93, 76 isolated units from CA1 pyramidal layer of Mouse 2, 3 and 4 respectively.

**Fig S12.**
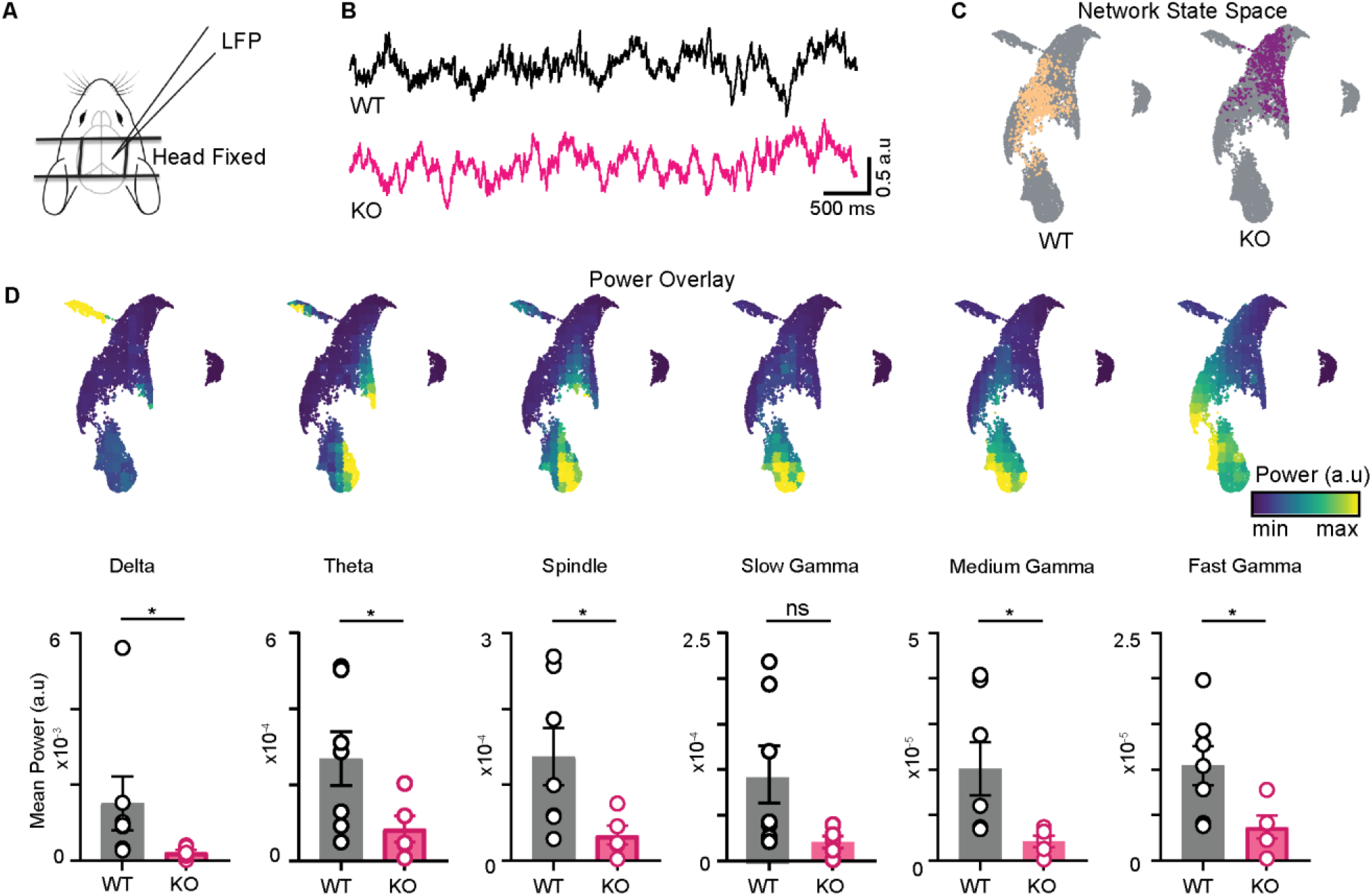
Altered organization of hippocampal oscillations in NLG3 KO mice A) Experimental paradigm: LFP recordings from hippocampal CA1 of head fixed mice at rest on a platform. B) Sample LFP traces obtained from WT (black) and KO (pink) mice. C) Combined network state space obtained by merging data from all trials and all animals (gray). Sample trial of WT and KO mice overlaid on combined network state space. D) Top: Power overlay on network state space for 6 oscillations of CA1. Bottom: Mean power comparison of individual bands. (WT vs KO, all power in arbitrary units) Delta (15*e* − 4 ± 7*e* − 4 *versus* 2*e* − 4 ± 7*e* − 5); Theta (26*e* − 5 ± 7*e* − 5 *versus* 8*e* − 5 ± 3*e* − 5); Spindle (13*e* − 5 ± 3*e* − 5 *versus* 3*e* − 5 ± 1*e* − 5); Slow Gamma (9*e* − 5 ± 3*e* − 5 *versus* 2*e* − 5 ± 6.6*e* − 6); Medium Gamma (2*e* − 5 ± 5.8*e* − 6 *versus* 4.2*e* − 6 ± 1.2*e* − 6); Fast Gamma (1*e* − 6 ± 2*e* − 6 *versus* 3.7*e* − 6 ± 1.2*e* − 6); Mann-Whitney test (All p<0.05 except for Slow Gamma)

**Fig S13.**
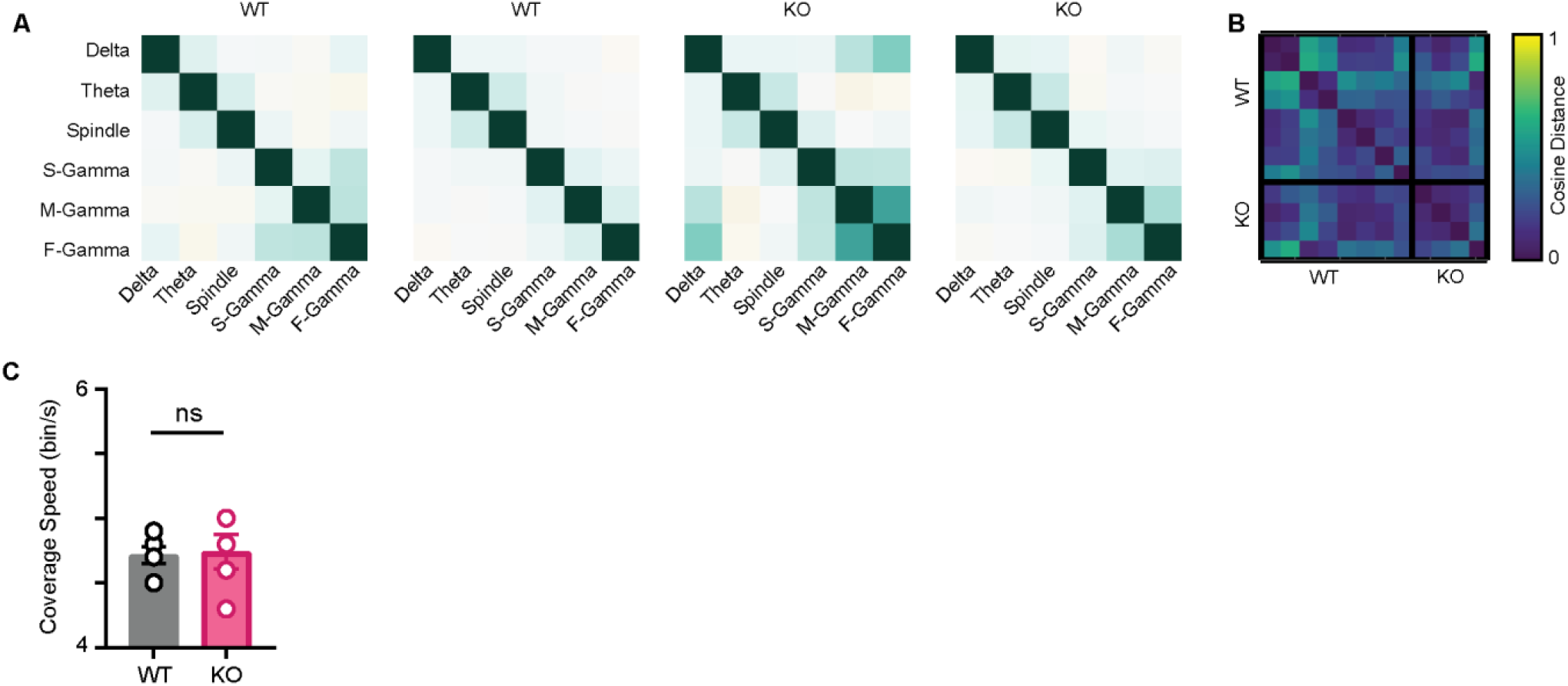
Characterization of state space in WT and NLG3 KO mice. A) Representative correlation matrix of oscillations from 2 trials of WT and KO mice. B) Pairwise cosine distance between correlation matrices from 7 trial of WT and 5 trials KO mice. Note the similarity among matrices across genotypes. C) Mean coverage speed for WT and KO mice (n = 7 WT trials and 5 NLG3 KO trials) (4.8 ± 0.08 *versus* 4.7 ± 0.1) p = 0.9, Mann-Whitney test

## Notes

### Competing Interest Statement

The authors have declared no competing interest.

